# A structure-based epitope tagging approach identifies vulnerable sites on the malarial P36-P52 protein complex for antibody-mediated neutralization of *Plasmodium* sporozoites

**DOI:** 10.64898/2026.03.16.712187

**Authors:** Samhita Das, Lien Boeykens, Manon Loubens, Carine Marinach, Sylvie Briquet, Line De Vocht, Isabel Pintelon, Jean-Pierre Timmermans, Yann G.-J. Sterckx, Olivier Silvie

**Affiliations:** Sorbonne Université, INSERM, CNRS, Centre d’Immunologie et des Maladies Infectieuses, Cimi-Paris, 75013 Paris, France; Laboratory of Medical Biochemistry and Infla-Med Centre of Excellence, University of Antwerp, Campus Drie Eiken, 2610 Wilrijk, Belgium; Laboratory of Cell Biology and Histology and the Antwerp Centre for Advanced Microscopy (ACAM), University of Antwerp, Campus Drie Eiken, 2610 Wilrijk, Belgium

**Author notes:** Contributed equally to the work. Contributed equally to the work (Samhita Das, Lien Boeykens).

**Keywords:** *Plasmodium*, malaria, sporozoites, 6-cys proteins, protein structure, neutralizing antibodies

## Abstract

Malaria is caused by apicomplexan parasites of the genus *Plasmodium*, which are transmitted through the bite of *Anopheles* mosquitoes that inject sporozoites (SPZs) into the skin. SPZs migrate to and infect the liver for an initial round of replication. SPZs and liver stages have long been considered as ideal targets for malaria vaccines. The main SPZ surface protein, the circumsporozoite protein (CSP), is the target of currently approved malaria vaccines and prophylactic antibody therapies. Studies in rodent malaria models have shown that anti-CSP antibodies exert their protective effect mainly in the skin, but some of the most potent anti-CSP monoclonal antibodies show additional protective effects in the vasculature and liver. Other SPZ proteins involved at different steps of the infection process may thus represent additional targets for antibody-mediated neutralization. Three 6-cysteine (6-Cys) domain proteins (P36, P52 and B9) play an essential role during SPZ invasion of hepatocytes, yet their molecular function and whether they can be targeted by neutralizing antibodies remains unknown. Here, to fill this gap, we combined an integrative structural biology approach with functional experiments in the *P. berghei* rodent malaria model. AlphaFold-based structural modeling followed by experimental validation via electron microscopy and small-angle X-ray scattering indicated that the P36-P52 heterodimer displays a head-to-tail architecture with an interaction interface that is largely conserved among *Plasmodium* species. The structural models supported the rational design of an epitope tagging approach, which, combined with neutralizing assays, revealed that antibodies against P36 and P52 can efficiently block invasion of hepatocytes by SPZs in culture conditions. The data show that the inhibitory activity of antibodies heavily depends on epitope position and revealed that antibody-exposed vulnerable sites lie on the membrane-distal side of the P36-P52 heterodimer. In contrast, antibodies targeting B9 had no inhibitory effect on SPZ invasion, irrespective of epitope positioning. These data show that the invasion step could be targeted by antibodies and indicate that the P36-P52 complex may be considered as a potential target for the development of next generation pre-erythrocytic malaria vaccines or therapeutic antibodies.

## Introduction

Malaria is caused by *Plasmodium*, a protozoan transmitted by female *Anopheles* mosquitoes. *P. falciparum* is the deadliest human-infective malaria parasite species, responsible for half a million deaths every year, mainly children under five in sub-Saharan Africa. *P. vivax* is widely distributed and the second causative agent of human malaria. *P. vivax* malaria is less severe than *P. falciparum*, but this species can persist in dormancy in infected individuals and cause malaria relapses weeks or months after the initial infection. Effective measures for malaria control (including the use of insecticide-treated bed nets and potent artemisinin-based antimalarial treatments) have resulted in a significant decrease in malaria incidence and mortality in many endemic areas, but progress in malaria control has recently stalled (WHO World Malaria Report 2025). Vaccines against *P. falciparum* and *P. vivax* would offer a safe and cost-effective way to protect large populations exposed to the malaria parasite in endemic areas.

Malaria begins with the inoculation of sporozoites (SPZs) into the host skin by infected *Anopheles* mosquitoes. The motile SPZs enter the blood stream and, upon reaching the liver, actively invade hepatocytes, where they differentiate into thousands of merozoites. Once released in the blood, merozoites invade and multiply inside erythrocytes, causing the malaria disease. Liver infection is an essential and clinically silent phase of the malaria life cycle, and constitutes an ideal target for a malaria vaccine. Blocking the parasite at the liver stage not only protects against the pathology but also abolishes parasite transmission to the mosquito vector, both associated with the subsequent blood stages.

The two currently approved malaria vaccines (RTS,S/AS01 and R21/MatrixM) both target the *P. falciparum* circumsporozoite protein (CSP), the main surface antigen of the extracellular SPZ stage. They both consist of a recombinant form of the *P. falciparum* CSP fused to the HBs antigen from Hepatitis B virus and adjuvanted in AS01 or Matrix M, respectively [1–4]. Protection induced by these vaccines is associated with CSP-specific antibodies [5]. The final results from a Phase III trial conducted in Africa have shown an overall reduction of 28-36% and 18-26% of clinical cases in RTS,S vaccinated young children and infants, respectively, depending on the immunization schedule [3,4]. R21 has shown higher efficacy in clinical trials, with protection up to 75% [1,2]. In addition to vaccines, immunotherapies based on monoclonal antibodies (mAbs) have recently emerged as a potential tool for malaria control [6]. In particular, clinical trials based on anti-CSP mAb therapies have established the proof of concept that antibody interventions can prevent malaria infection [7–10]. Mechanistic studies in rodent malaria models have shown that anti-CSP antibodies exert their neutralizing activity mainly in the skin, through inhibition of gliding motility and/or direct killing of the parasite [11–13]. Recent studies provided *in vivo* evidence that the most potent anti-CSP mAbs neutralize SPZs not only in the skin but also in the vasculature and liver, pointing at these processes as important targets to achieve optimal protection [13,14]. This suggests that other SPZ antigens in addition to CSP could be valuable targets for vaccines and mAbs, including proteins mediating parasite host cell invasion in the liver.

In previous studies, we have characterized host entry pathways used by human and rodent parasites to infect hepatocytes [15,16]. We further identified a SPZ protein, the 6-cysteine domain (6-Cys) protein P36, as a key determinant of host cell receptor usage in rodent malaria parasites, establishing for the first time a functional link between SPZ and host cell entry factors [16]. The 6-Cys fold features a combination of parallel and anti-parallel β-sheets and the presence of six conserved cysteines that form three intramolecular disulfides [17]. There are two 6-Cys domain types (called A and B) with slightly different topologies [18,19] and the 6-Cys domains of different *Plasmodium* 6-Cys proteins are rather poorly conserved at the amino acid sequence level despite a high structural similarity (which is a typical feature of parasite surface antigens [20]). The 6-Cys fold resembles the topology of ephrins, a family of surface-exposed, membrane-anchored proteins mainly involved in cell-cell interactions in multicellular organisms. The 6-Cys domain is thought to have been obtained from the vertebrate host by an ancestral apicomplexan parasite through horizontal gene transfer. Over the course of time, several apicomplexans have tailored the adhesive properties of the acquired 6-Cys domain to their own benefit [21,22]. The outcome is the occurrence of parasite 6-Cys proteins containing canonical 6-Cys domains and/or “degenerate” ephrin-like domains containing only 2 or 4 cysteines (2-Cys and 4-Cys). *Plasmodium* 6-Cys proteins are expressed throughout the parasite’s life cycle, where they are involved in the biology of extracellular invasive and intracellular replicative stages [23–25]. A recurring theme is that simultaneously expressed 6-Cys proteins form complexes at the parasite surface (typically heterodimers), in which one partner secures membrane attachment through a C-terminal glycosylphosphatidylinositol (GPI) anchor. This has been confirmed for 6-Cys protein complexes P48/45-P230 (gametocyte) [26,27], P12-P41 (merozoite) [28,29], and P36-P52 (SPZ) [30].

Gene knockout studies in *P. falciparum*, *P. berghei* and *P. yoelii* established that both P36 and P52, as well as a third member of the 6-Cys family, the propeller domain-containing B9 protein, are required for productive invasion of hepatocytes [16,31–34]. All three proteins are contained in the SPZ micronemes, and are secreted upon parasite activation [30–32]. However, their function is still unknown at the molecular level. Given their role during SPZ invasion, P36, P52 and B9 could represent potential vaccine targets. However, whether these proteins can be targeted by neutralizing antibodies remains unknown. Here, we use a highly interdisciplinary strategy in which we employ experimentally validated AlphaFold [36,37] models of the P36-P52 heterodimer complex to rationally design an epitope tagging approach in the *P. berghei* rodent malaria model to address whether antibodies targeting P36, P52 or B9 can inhibit infection of hepatocytes by SPZs.

## Results

### The P36-P52 heterodimer displays a head-to-tail architecture with an interaction interface that is largely conserved among *Plasmodium* species

Apart from evidence supporting the existence of a P36-P52 heterodimer [30], its structural details are yet to be presented. Furthermore, it is not clear whether other SPZ 6-Cys proteins such as B9 are involved in the formation of larger 6-Cys complexes [32]. To answer these questions, we performed structure prediction of the P36-P52 heterodimer using AlphaFold-Multimer [35,36]. The structures of P36-P52 complexes from *P. falciparum, P. vivax, P. berghei* and *P. yoelii* could be predicted with relatively high confidence as judged from various metrics (pTM, ipTM, pLDDT, PAE, and zDOPE; **Figure 1**). The P36-P52 heterodimers display an antiparallel “head-to-tail” arrangement in which the P52 N-terminal and C-terminal domains (P52-6D1 and P52-6D2, respectively) interact with the P36 C-terminal and N-terminal 6-Cys domains (P36-6D2 and P36-6D1, respectively), respectively (**Figure 2**). While there are also interactions between the P36 and P52 C-terminal domains (P36-6D2 and P52-6D2), there are no contacts between the N-terminal domains (P36-6D1 and P52-6D1). Given that P52 is predicted to contain a C-terminal GPI anchor, this would imply that the P36-6D1 and P52-6D2 domains are in relative proximity to the parasite membrane, whereas the P36-6D2 and P52-6D1 domains are membrane-distal (**Figure 2**). For all species, the interaction interfaces on the 6D1 and 6D2 domains of both proteins are relatively flat, which is in contrast with the P12-P41 complex [19,29]. Further inspection reveals that the contacting surface areas are extensive (∼1600 to 1800 Å^2^) (**Table 1**) and consist of both hydrophilic and hydrophobic interactions. An *in silico* analysis of the interaction strength suggests that the P36-P52 heterodimers could be high-affinity complexes (**Table 1**), although this remains to be experimentally confirmed. Interestingly, the heterodimer interface appears to be relatively well conserved across *Plasmodium* species. This conservation feature becomes apparent upon i) coloring of the heterodimer surface interface by hydrophobicity (**Figure 2**, middle sections), ii) visualizing the interface via an interaction heat map (**Figure 2**, bottom sections), and iii) analysis of a multiple sequence alignment (**Supplementary Figure S1**). While the first two reveal the emergence of a conserved interaction pattern, the latter reveals 54.3% and 61.3% pairwise identities for the P36 and P52 interface residues, respectively (compared to overall pairwise identities of 52.0% and 40.3%, respectively).

**Figure 1.**
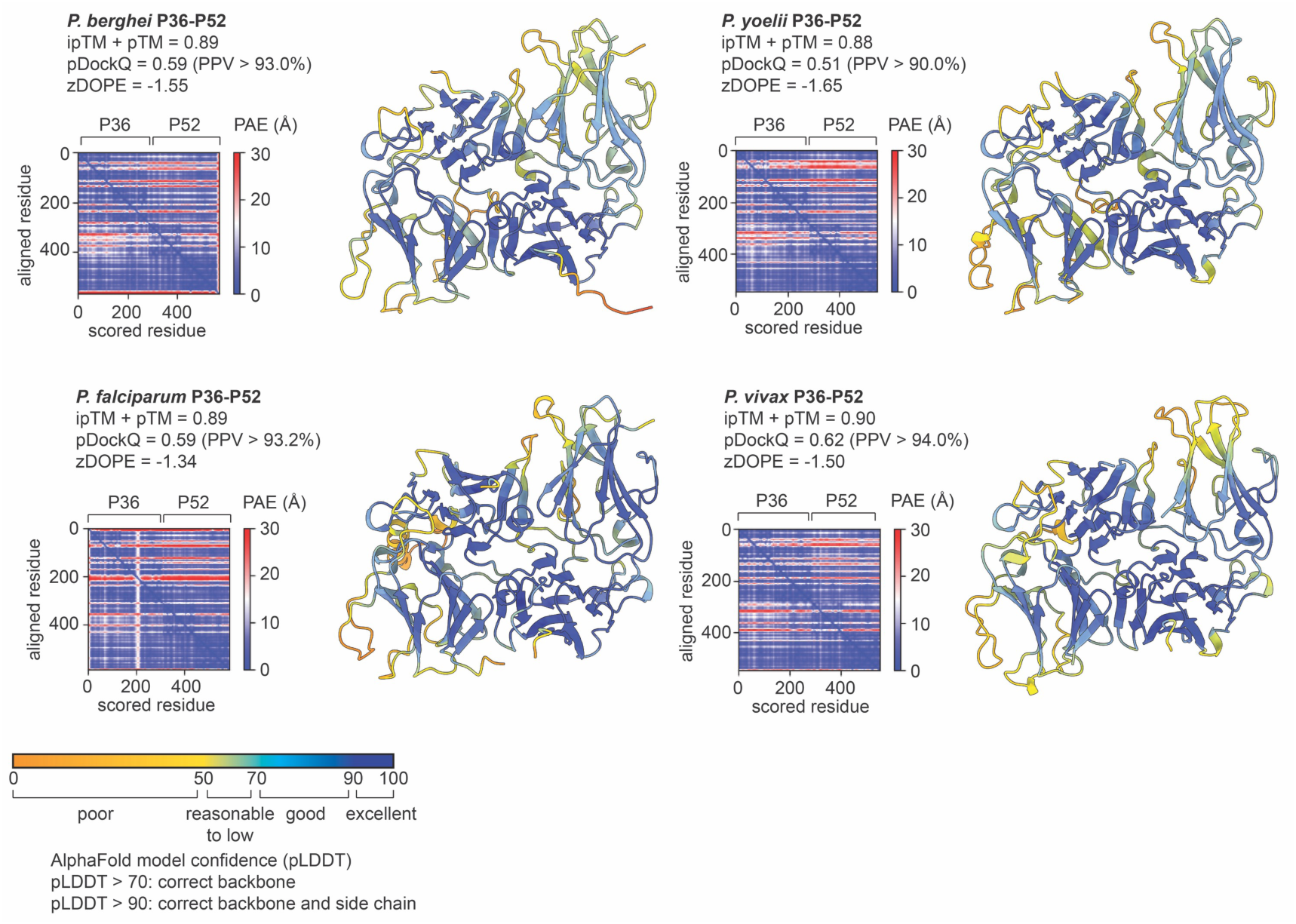
Overview of the AlphaFold prediction models for the P36-P52 heterodimers and associated quality metrics. The models are displayed as cartoon representations and are colored according to the predicted local distance difference test (pLDDT) score, which reflects (local) model quality as indicated by the legend at the bottom. For all structures, the predicted aligned error (PAE), the normalized discrete optimized protein energy (zDOPE), the pDockQ and AlphaFold-Multimer model confidence (0.8*ipTM + 0.2*pTM) are also shown. The PAE provides a distance error for every residue pair, and is calculated for each residue x (scored residue) when the predicted and true structures are aligned on residue y (aligned residue). A zDOPE < -1 indicates that the distribution of atom pair distances in the model resembles that found in a large sample of known protein structures and that at least 80% of the model’s Cα atoms are within 3.5 Å of their correct positions. The pTM (score between 0 and 1) provides a measure of similarity between two protein structures (in this case, the predicted and unknown true structure) over all residues and thus reports on the accuracy of prediction within a single chain. The interface pTM (ipTM, score between 0 and 1) provides a measure of similarity between two protein structures (in this case, the predicted and unknown true structure) over only interfacing residues and thus reports on the accuracy of prediction for a complex. The pDockQ score (between 0 and 1) is another confidence metric for protein complexes that considers the number of interfacing residues and their pLDDT scores. Determination of the pDockQ score can be associated to a positive predictive value (PPV), which provides an estimate for the probability that the solution is a true positive.

**Figure 2.**
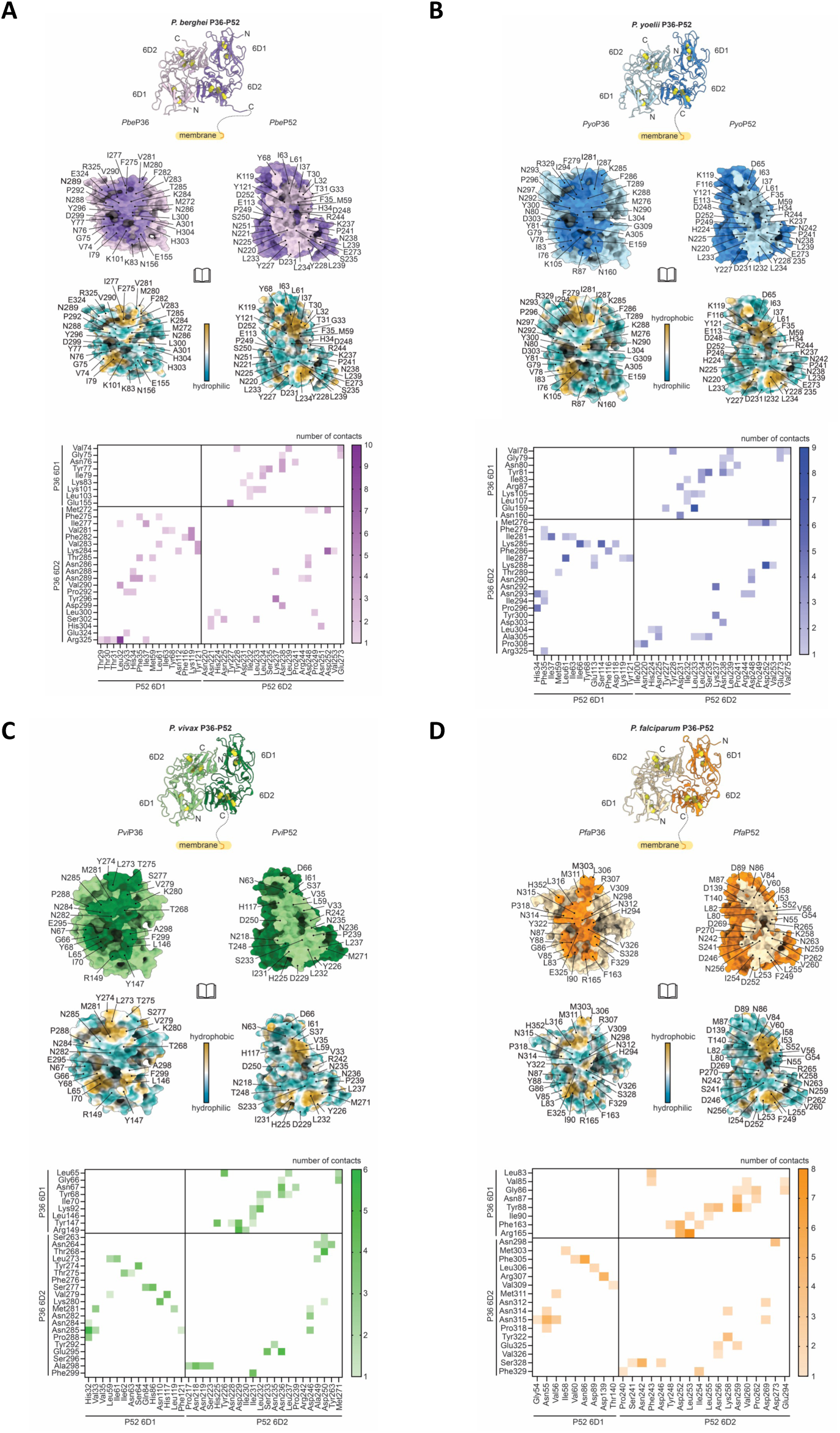
Analysis of the interactions at the P36-P52 heterodimer interfaces. The results are shown for *P. berghei*, *P. yoelii*, *P. vivax*, and *P. falciparum* P36-P52 (panels **A**, **B**, **C**, and **D**, respectively). In all panels, the top sections show a ribbon representation of the P36-P52 heterodimers considering their membrane attachment via P52’s GPI anchor. For each complex, P36 and P52 are colored in light and dark shades, respectively, of purple (*P. berghei*), blue (*P. yoelii*), green (*P. vivax*) and orange (*P. falciparum*). Disulfides are shown as spheres and colored by heteroatom. The middle sections display two surface representations of the P36 (left) and P52 (right) interaction interfaces for each complex. In the first surface representation, the surfaces of P36 and P52 are color-coded as in the top panel and the residues involved in complex formation bear the color of the interacting partner (dark color shade on the P36 surface and light color shade on the P52 surface). The second representation shows the P36 and P52 surfaces in the same orientation colored according to their molecular hydrophobicity potential. The bottom sections display interaction heat maps, which, together with the analysis displayed in the middle sections and when compared to each other, underline the conserved feature of the P36-P52 interaction interface within *Plasmodium* species.

**Table 1.**
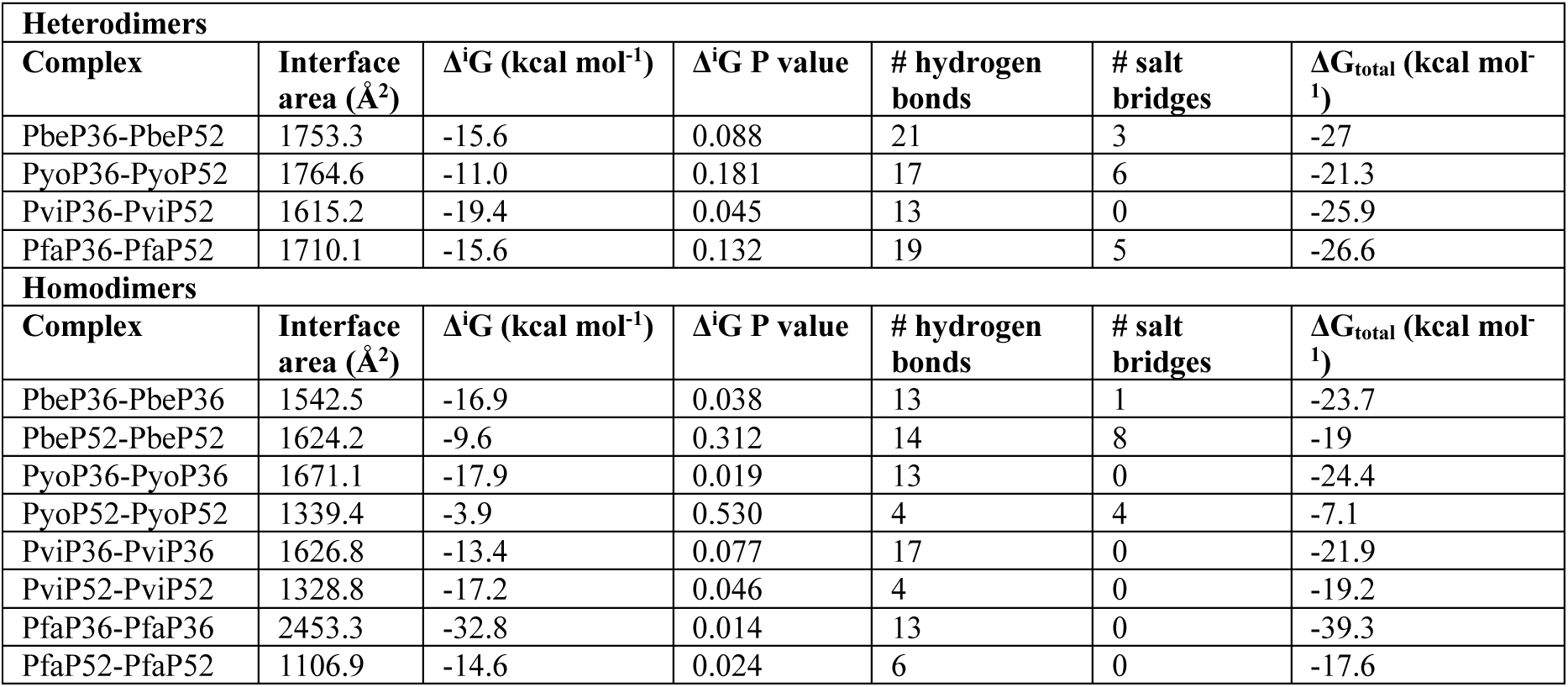
PISA interface analysis of the P36-P52 complexes.

| **Heterodimers** |  |  |  |  |  |  |
| --- | --- | --- | --- | --- | --- | --- |
| **Complex** | **Interface area (Å<sup>2</sup>)** | **Δ<sup>i</sup>G (kcal mol<sup>-1</sup>)** | **Δ<sup>i</sup>G P value** | **# hydrogen bonds** | **# salt bridges** | **ΔG<sub>total</sub> (kcal mol<sup>-1</sup>)** |
| PbeP36-PbeP52 | 1753.3 | -15.6 | 0.088 | 21 | 3 | -27 |
| PyoP36-PyoP52 | 1764.6 | -11.0 | 0.181 | 17 | 6 | -21.3 |
| PviP36-PviP52 | 1615.2 | -19.4 | 0.045 | 13 | 0 | -25.9 |
| PfaP36-PfaP52 | 1710.1 | -15.6 | 0.132 | 19 | 5 | -26.6 |
| **Homodimers** |  |  |  |  |  |  |
| **Complex** | **Interface area (Å<sup>2</sup>)** | **Δ<sup>i</sup>G (kcal mol<sup>-1</sup>)** | **Δ<sup>i</sup>G P value** | **# hydrogen bonds** | **# salt bridges** | **ΔG<sub>total</sub> (kcal mol<sup>-1</sup>)** |
| PbeP36-PbeP36 | 1542.5 | -16.9 | 0.038 | 13 | 1 | -23.7 |
| PbeP52-PbeP52 | 1624.2 | -9.6 | 0.312 | 14 | 8 | -19 |
| PyoP36-PyoP36 | 1671.1 | -17.9 | 0.019 | 13 | 0 | -24.4 |
| PyoP52-PyoP52 | 1339.4 | -3.9 | 0.530 | 4 | 4 | -7.1 |
| PviP36-PviP36 | 1626.8 | -13.4 | 0.077 | 17 | 0 | -21.9 |
| PviP52-PviP52 | 1328.8 | -17.2 | 0.046 | 4 | 0 | -19.2 |
| PfaP36-PfaP36 | 2453.3 | -32.8 | 0.014 | 13 | 0 | -39.3 |
| PfaP52-PfaP52 | 1106.9 | -14.6 | 0.024 | 6 | 0 | -17.6 |

### In-solution structural studies experimentally confirm the head-to-tail architecture of the P36-P52 heterodimer

To experimentally confirm the P36-P52 heterodimer AlphaFold model, we attempted to recombinantly produce P36 and P52 as separate proteins. Unfortunately, despite various trials (including variations in heterologous expression systems) this proved to be troublesome in our hands. Typical production and purification campaigns were plagued by relatively low yields (averages of ∼100 µg per liter of culture), problems in reproducibility, recombinant production of the target protein in the insoluble fraction, or no recombinant production at all depending on the construct. The best results were obtained for *P. falciparum* P52 in insect cells (Sf21) with yields of ∼145 µg per liter of culture. Interestingly, besides the occurrence of *P. falciparum* P52 monomer fractions, we noticed a significant propensity for the protein to form oligomers (**Supplementary Figure S2**). Our observations appear to be consistent with those published by others for *P. yoelii* P52 [37,38]. The formation of P36-P36 and P52-P52 homodimers might indeed be possible given that the structural analyses for the P36-P52 heterodimer indicate that the interaction interfaces on the 6D1 and 6D2 domains of both proteins are relatively flat (**Figure 2**). Intrigued by these observations, we employed AlphaFold to investigate this hypothesis. Interestingly, AlphaFold modeling supports the occurrence of such homodimers as judged from various quality metrics (pTM, ipTM, pLDDT, PAE, and zDOPE; **Supplementary Figure S3**). A possible biological relevance for these homodimers (if any) remains enigmatic.

As the recombinant production of P36 and P52 as individual proteins proved to be highly cumbersome and the existence of P36-P36 and P52-P52 homodimers might preclude successful reconstitution of the P36-P52 heterodimer *in vitro*, we hypothesized that P36 and P52 may need to be co-produced. To increase the chance of obtaining correctly folded P36-P52 heterodimers, we generated fusion constructs in which we connected the P52 C-terminus to the P36 N-terminus through a (GGGGS)4 linker given that the AF models strongly suggested a head-to-tail arrangement. Recombinant production in Sf9 insect cells could be obtained for the *P. falciparum* P52-P36 fusion protein, although the yields after purification remained rather low (∼240 µg per liter of insect cell culture). In addition, our sample preparations often still contain an insect cell contaminant, which was identified to be Sf9 cathepsin L (**Supplementary Figure S4**). Nonetheless, the in-solution structure of the *P. falciparum* P52-P36 fusion protein could still be probed via negative stain electron microscopy (EM) and small-angle X-ray scattering (SAXS) (**Figure 3**). Both data sets concur with the head-to-tail architecture for the *P. falciparum* P36-P52 heterodimer, thereby experimentally validating the AlphaFold prediction models. Given the strong conservation of the P36-P52 interaction interface within *Plasmodium* species, this architecture is very likely to be the same for the *P. berghei*, *P. yoelii*, and *P. vivax* P36-P52 complexes. The low yields and issues with the Sf9 cathepsin L contamination, currently make further structural and functional studies with this construct difficult. We are currently optimizing the production and purification conditions. In the meantime, we opted to continue our work with an epitope tagging approach, rationally designed on the experimentally validated head-to-tail architecture of the P36-P52 heterodimer.

**Figure 3.**
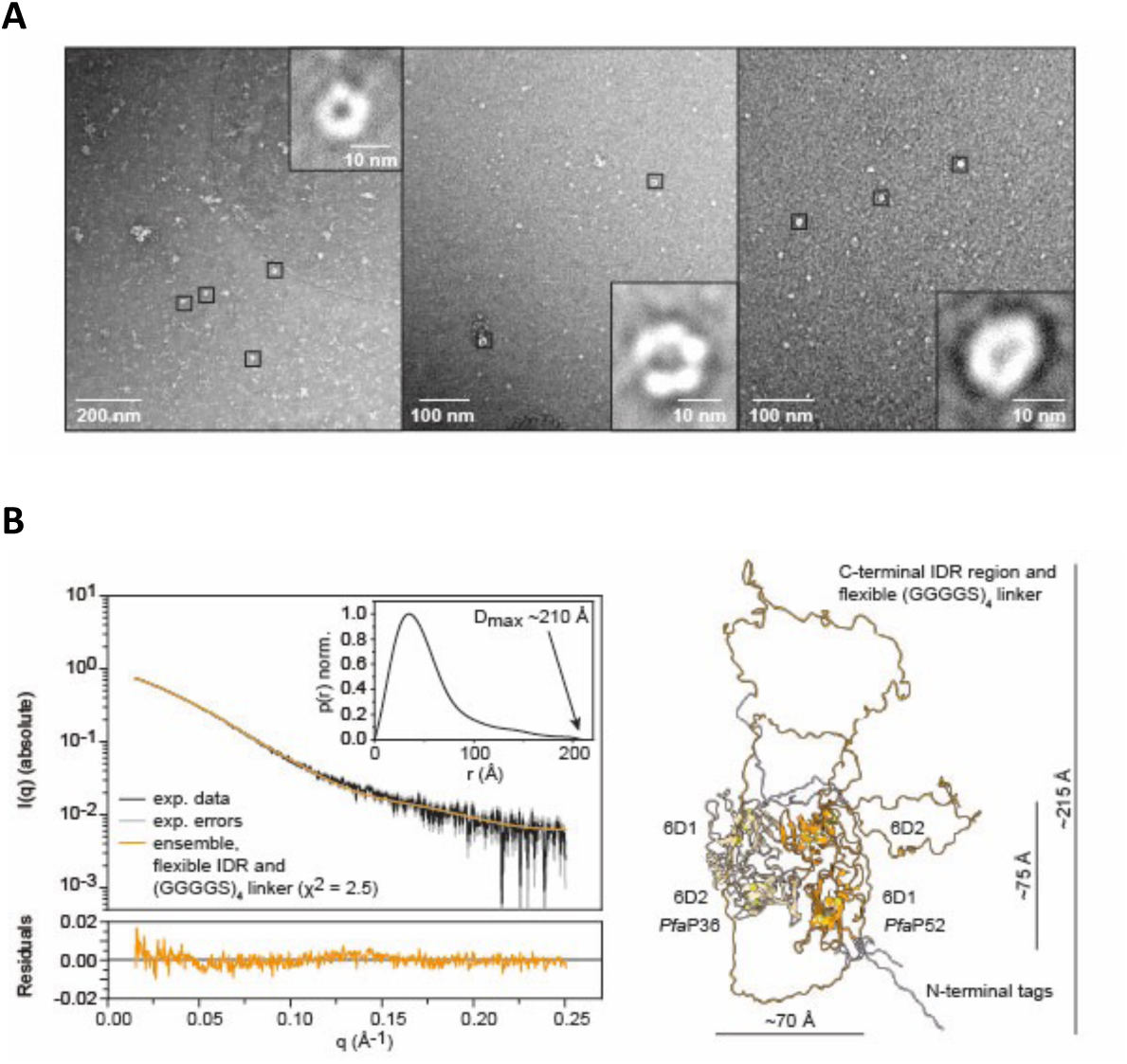
Experimental validation of the P36-P52 AlphaFold prediction model. **A**. Negative stain EM micrographs collected for the *P. falciparum* P52-P36 fusion protein, which show the occurrence of particles with shapes and dimensions consistent with the AlphaFold prediction model (indicated by the black squares). **B**. SEC-SAXS analysis on the *P. falciparum* P52-P36 fusion protein fraction (see **Supplementary Figure S4**). The left graph shows the experimental data (black), error margins (gray), and the fits of a conformational ensemble considering (GGGGS)4 linker flexibility (orange) to the data. The inset shows a normalized probability distance distribution function (PDDF) obtained from the experimental SAXS data, with the approximate Dmax value indicated. The PDDF bears all hallmarks of a globular particle (the P36-P52 heterodimer) with a highly flexible moiety (the (GGGGS)4 linker). The right panel shows the conformational ensemble obtained for the *P. falciparum* P52-P36 fusion protein.

### Antibodies targeting the membrane-distal domains of the P36-P52 complex inhibit SPZ invasion *in vitro*

Failure to produce large quantities of recombinantly expressed P36 and/or P52 was a major limitation for the generation of specific antibodies and the assessment of antibody neutralizing activity. To bypass the need for recombinant parasite antigens, we designed a novel strategy based on structure-guided genetic epitope tagging and the use of anti-tag mAbs in functional assays (**Figure 4A**). We implemented this strategy in the *P. berghei* model to address whether antibodies targeting tagged versions of P36 and P52 can inhibit SPZ infectivity. As established in the section above, AlphaFold predicts the formation of a P36-P52 heterodimer in a head-to-tail orientation, with the N-terminal and C-terminal domains of P36 and P52, respectively, being membrane-proximal and the C-terminal and N-terminal regions of P36 and P52, respectively, being membrane-distal. We reasoned that the position of the epitope in the protein may affect the capacity of antibodies to neutralize its function. To test this hypothesis, we introduced an epitope tag either at the N-terminus or at the C-terminus of P36 and P52 (**Figure 4B**).

**Figure 4.**
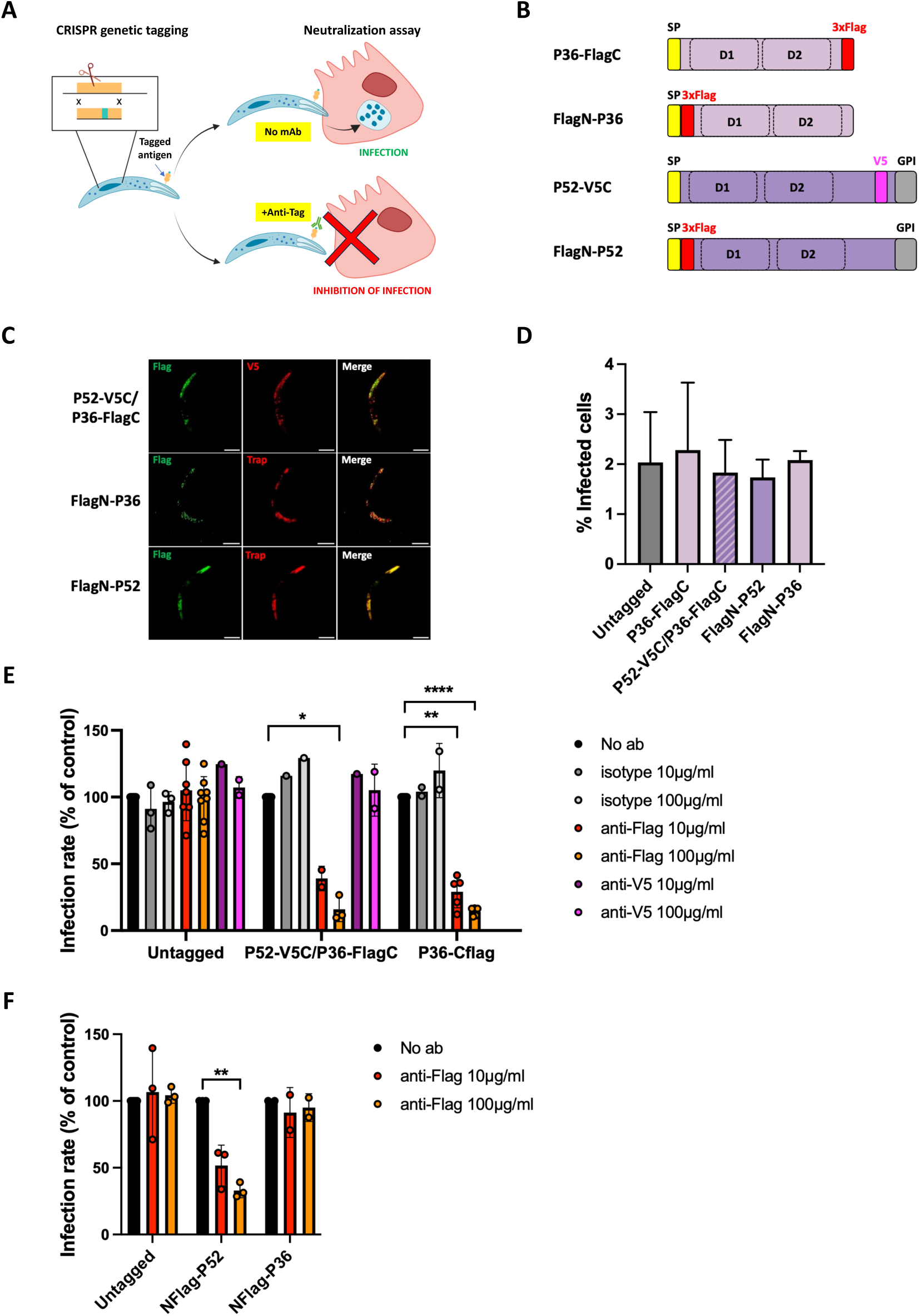
SPZ invasion inhibition assay using anti-tag antibodies. **A**. Overview of the functional assay to test the neutralizing activity of anti-tag antibodies on transgenic SPZs expressing a tagged version of a protein of interest. **B**. Schematic representation of the different P36/P52 constructs tested in this study. **C**. Expansion microscopy images of P52-V5C/P36-FlagC, FlagN-P36 and FlagN-P52 sporozoites using anti-Flag antibodies (green), anti-V5 or anti-TRAP antibodies (red). Scale bar, 10 μm. **D.** Infectivity of transgenic SPZs to HepG2 cells was analyzed by flow cytometry 24 h post-infection. The data represent the % of GFP-positive cells (mean +/- SEM of independent experiments; each symbol represents the mean value in one experiment). **E**. HepG2 cells were infected with control untagged parasites, P52-V5C/P36-FlagC or P36-FlagC GFP-expressing *P. berghei* SPZs, in the presence of anti-Flag, anti-V5, or isotype control antibodies. Infected cells were quantified 24 h post-infection by flow cytometry. The data represent the % of GFP-positive cells, expressed as the % of the control without antibody (mean +/- SEM of independent experiments; each symbol represents the mean value of one experiment). **F.** HepG2 cells were infected with control untagged parasites, FlagN-P36 or FlagN-P52 GFP-expressing *P. berghei* SPZs, in the presence of anti-Flag antibodies. Infected cells were quantified 24 h post-infection by flow cytometry. The data represent the % of GFP-positive cells, expressed as the % of the control without antibody (mean +/- SEM of independent experiments; each symbol represents the mean value of one experiment). *, p<0.05; **, p<0.01; ****, p<0.0001 (two-tailed ratio paired t test).

Parasite lines expressing C-terminally tagged P52 and/or P36 were generated through genetic complementation of drug selectable marker-free GFP-expressing Δ*p52p36* and Δ*p36 P. berghei* ANKA parasites [16]. In a previous study, we generated parasites expressing a tagged version of B9 (here referred to as B9-FlagC), where a 3xFlag epitope was introduced after the putative omega site toward the C-terminus [32]. We used a similar strategy to introduce a V5 epitope in P52, which like B9 is also predicted to be GPI-anchored. In the case of P36, which does not contain any membrane associated domain at its C-terminus, a 3xFlag epitope was added at the end of the protein just upstream of the STOP codon. Δ*p36* parasites were complemented by double cross-over homologous recombination with a construct encoding a P36-3xFlag cassette, to generate P36-FlagC parasites, carrying a 3xFlag epitope tag at the C-terminus of P36 (**Supplementary Figure S5**). Δ*p52p36* parasites were complemented by double cross-over homologous recombination with a construct encoding a double P52-V5/P36-3xFlag cassette, resulting in the P52-V5C/P36-FlagC parasites (**Supplementary Figure S5**). Following transfection, recombinant parasites were selected with pyrimethamine and genotyped by PCR (**Supplementary Figure S5**). We used CRISPR-Cas9 [39] to generate two additional parasite lines, referred to as FlagN-P36 and FlagN-P52, expressing N-terminally tagged version of P36 and P52, respectively (**Figure 4B**). A 3xFlag epitope was inserted after the signal peptide, just upstream of the first 6-Cys domain in each of the proteins. Following transfection of PbCasDiCre-GFP acceptor parasites [40], genetically modified FlagN-P36 and FlagN-P52 parasites were obtained and verified by PCR and Sanger sequencing of the amplicons.

Genetically modified parasites were then transmitted to *Anopheles stephensi* mosquitoes, and SPZs were collected from the salivary glands of infected mosquitoes 3 weeks after transmission. We confirmed by immunofluorescence assays (IFA) that the tagged proteins were readily detected in salivary gland SPZs (**Figure 4C**). SPZs from genetically complemented parasites invaded HepG2 cells as efficiently as parental parasites, as evidenced by flow cytometry (**Figure 4D**), showing that introduction of the tags had no deleterious effect on the function of P52 and P36.

Next, we tested whether antibodies against V5 or Flag tags could inhibit invasion of HepG2 cells by SPZs, starting with P36-FlagC and P52-V5C/P36-FlagC parasite lines. As a control, we used PbGFP parasites, which express the original untagged version of the 6-Cys proteins. HepG2 cell cultures were incubated with SPZs in the presence of increasing concentrations of anti-tag antibodies, for 3 hours, then washed and further incubated for an additional 24-48 hours. Productive invasion was then quantified by flow cytometry, based on the percentage of infected GFP-positive cells. Anti-V5 and anti-Flag antibodies had no effect on the control PbGFP SPZ infectivity, ruling out any non-specific toxic effect of the antibody formulations (**Figure 4E**). Remarkably, anti-Flag antibodies inhibited invasion by SPZs expressing P36-FlagC, in a dose-dependent manner, while control antibodies had no effect (**Figure 4E**). The neutralizing activity of anti-Flag antibodies was observed with both the P52-V5C/P36-FlagC and P36-FlagC parasite lines. In sharp contrast, anti-V5 antibodies did not inhibit P52-V5/P36-FlagC SPZ invasion in culture conditions, even at the highest concentration tested (100 μg/mL) (**Figure 4E**).

FlagN-P36 and FlagN-P52 SPZs were also tested in the functional assay, along with untagged parental PbCasDiCre-GFP SPZ. As expected, the anti-Flag antibody had no effect on the untagged parasites used as a control (**Figure 4F**). In sharp contrast, anti-Flag antibodies inhibited FlagN-P52 SPZ invasion of HepG2 cells, in a dose-dependent manner (**Figure 4F**). Although this effect was less pronounced than with P36-FlagC parasites, it shows that P52, like P36, is accessible to neutralizing antibodies. Interestingly, the anti-Flag antibody had no effect on FlagN-P36 SPZs (**Figure 4F**), reminiscent of our observations with the P52-V5C parasites and anti-V5 antibodies. Altogether, these data show that the position of the epitope affects the neutralizing activity of anti-tag antibodies, and suggest that the C-terminal 6-Cys domain (6D2) of P36 and the N-terminal 6-Cys domain (6D1) of P52 are vulnerable regions in the complex that can be targeted by neutralizing antibodies.

### Antibodies targeting B9 do not inhibit SPZ invasion

We used the same functional assay as described above to assess whether antibodies can neutralize SPZs by targeting the 6-Cys protein B9. We used B9-FlagC parasites [32], and generated an additional FlagN-B9 line with a 3xFlag epitope introduced at the N-terminus of the protein, immediately upstream of the beta propeller domain (**Figure 5A** and **Supplementary Figure S5**). B9-FlagC and FlagN-B9 parasites were transmitted to mosquitoes to produce SPZs and we verified expression of Flag-tagged B9 by immunofluorescence (**Figure 5B**). FlagN-B9 SPZs were fully infective in cell culture conditions, similar to B9-FlagC parasites (**Figure 5C**). We then tested the neutralizing activity of anti-Flag antibodies on FlagN-B9 and B9-FlagC, using the same functional assay as described above. Unlike for P36 and P52, anti-Flag antibodies failed to inhibit FlagN-B9 and B9-FlagC SPZs, even at the highest concentration tested (**Figure 5D**). These observations indicate that either B9 is not accessible to antibody-mediated neutralization, or that the N-terminus and C-terminus of the protein are not vulnerable regions.

**Figure 5.**
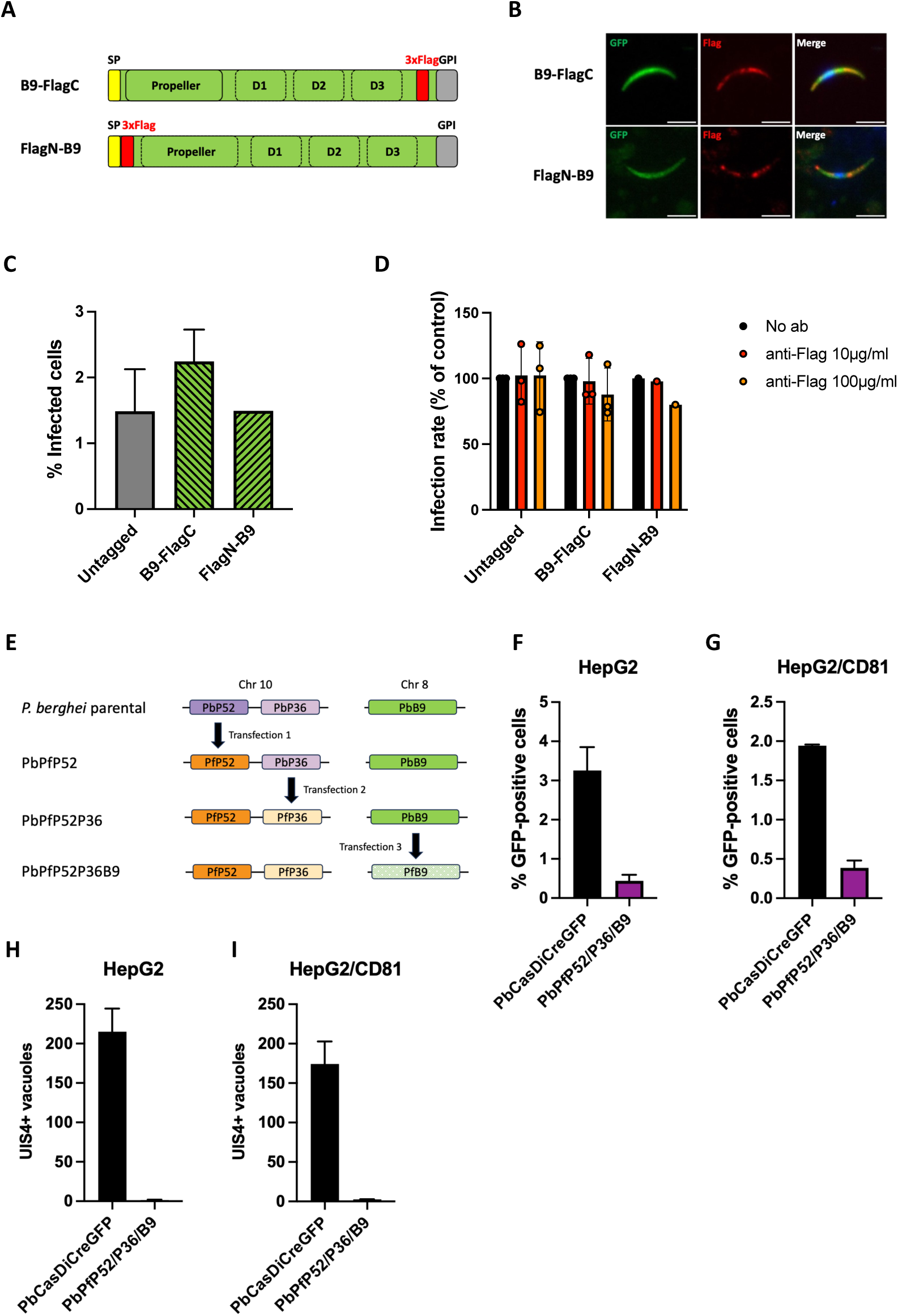
Antibodies targeting B9 do not inhibit SPZ invasion. **A**. Schematic representation of the B9 constructs tested in this study. **B**. Immunofluorescence analysis of GFP-expressing B9-FlagC and FlagN-B9 parasites using anti-Flag antibodies (red). Nuclei were stained with Hoechst 33342 (blue). Scale bar, 5 μm. **C**. Infectivity of untagged control parasites, FlagN-B9 and B9-FlagC SPZs to HepG2 cells was analyzed by flow cytometry 24 h post-infection. The data represent the % of GFP-positive cells (mean +/- SEM of independent experiments). **D**. HepG2 cells were infected with control untagged parasites, B9-FlagC or FlagN-B9 GFP-expressing *P. berghei* SPZs, in the presence of anti-Flag antibodies. Infected cells were quantified 24 h post-infection by flow cytometry. The data represent the % of GFP-positive cells, expressed as the % of the control without antibody (mean +/- SEM of independent experiments; each symbol represents the mean value of one experiment). **E**. CRISPR generation of PbPfP36P52B9 parasites, in three successive steps. **F-G**. Quantification of invaded HepG2 (F) and HepG2/CD81 (G) cells analyzed by flow cytometry 24 h post-infection with PbCasDiCreGFP (parental) or PbPfP52P36B9 salivary gland SPZs. The results shown are the mean +/- SEM of two independent experiments. The low level of invaded cells likely corresponds to the cell traversal events (not productive invasion). **H-I**. Quantification of UIS4-labelled exo-erythrocytic forms (EEFs) in HepG2 (H) and HepG2/CD81 (I) cells, as determined by fluorescence microscopy 48h post-infection. The results shown are the mean +/- SEM of two independent experiments.

### Absence of evidence supporting the existence of a functional complex between P36/P52 and B9

Previous inter-species genetic complementation studies showed that substitution of *P. berghei* P36 and P52 by *P. yoelii* homologous proteins maintained SPZ infectivity, while *P. falciparum* and *P. vivax* P36 and P52 were not functional in *P. berghei* [106]. Similarly, *P. berghei* B9 could be replaced by *P. yoelii* but not by *P. falciparum* B9 propeller [107]. Given the similar phenotype of P36/P52 and B9 gene deletion mutants, we hypothesized that the three 6-Cys proteins might be part of a functional tripartite complex required for SPZ invasion. Concordantly, heterologous expression of *P. berghei* proteins in mammalian cells showed an interaction between the propeller domain of B9 and either P36 or P52, although no such interaction could be detected in SPZ extracts by co-immunoprecipitation of B9 and mass spectrometry [107].

We first turned to AlphaFold to predict the structure of a putative P36-P52-B9 complex. Unfortunately, such complex could not be predicted as evidenced by the poor metrics (**Supplementary Figure S6**). However, it’s not because powerful algorithms such as AlphaFold cannot predict the structure, that such a complex may not exist. Hence, as an alternative approach to interrogate the existence of a functional tripartite P36-P52-B9 complex, we generated a *P. berghei* transgenic line expressing *P. falciparum* P36, P52 and B9, using CRISPR (**Figure 5E**). PbCasDiCre-GFP parasites were genetically modified sequentially to replace *P. berghei* P52 by *P. falciparum* P52, *P. berghei* P36 by *P. falciparum* P36, and *P. berghei* B9 by *P. falciparum* B9, resulting in the generation of a triple falciparumized parasite line PbPfP52P36B9 (**Supplementary Figure S7**). PbPfP52P36B9 parasites were obtained after three successive transfections and were genotyped by PCR to confirm the desired recombinations (**Supplementary Figure S7**). After transmission to mosquitoes, PbPfP52P36B9 SPZs were collected and analyzed *in vitro* in HepG2 and HepG2/CD81 cell cultures. PbPfP52P36B9 parasites showed very low levels of cellular invasion by flow cytometry in both cell types (**Figure 5F-G**), and no EEFs were seen after staining with UIS4, a marker for the parasitophorous vacuole membrane [41], irrespective of CD81 expression (**Figure 5H-I**). Hence, PbPfP52P36B9 fail to productively invade hepatocytic cells, reproducing a similar phenotype as observed before with P36-, P52- or B9-deficient SPZs and with parasites separately complemented with *P. falciparum* P36/P52 or *P. falciparum* B9 [16,32]. Altogether, these data do not support the existence of a functional P36/P52/B9 complex, and strongly suggest that other factors are involved to mediate SPZ invasion of hepatocytes.

## Discussion

Most studies of SPZ targeting by antibodies have focused on CSP. These studies have clearly established that anti-CSP antibodies exert their neutralizing effect mainly in the skin, where they can immobilize the parasite before it can migrate to the liver [11,13,42]. Anti-CSP antibodies are also able to exert direct cytotoxicity on SPZs [11,13], through a mechanism that relies on pore-forming proteins secreted by the parasite [11]. Nevertheless, recent work using human anti-CSP mAbs and transgenic *P. berghei* parasites expressing *P. falciparum* CSP has shown that the most potent mAbs have an additional effect in the vasculature and in the liver, and can also affect post-invasion liver stage development [13,14]. These observations suggest that targeting different steps of the SPZ journey, including host cell invasion in the liver, could be the most effective strategy to achieve optimal protection. P36 and P52 could be potential targets for such a combined approach.

Based on the encouraging report that AlphaFold can correctly predict heterodimer 6-Cys protein complexes (as demonstrated for the *P. falciparum* P12-P41 complex [17]), we used it here to predict the structures of the *P. falciparum*, *P. berghei*, *P. yoelii* and *P. vivax* P36-P52 heterodimers. These could all be predicted with high confidence and consistently displayed an antiparallel “head-to-tail” arrangement (as observed for *P. falciparum* P12-P41 [17,19,29]) in which the P36-6D1 and P52-6D2 domains are located membrane-proximal and the P36-6D2 and P52-6D1 domains membrane-distal. The “head-to-tail” arrangement was experimentally confirmed for *P. falciparum* P36-P52 through negative stain EM and SAXS on a recombinantly obtained fusion protein. *In silico* analysis of the interaction interfaces suggests that these are high affinity complexes and that they are well conserved across *Plasmodium* species. This most probably explains why “mixing and matching” of P36 and P52 proteins from different *Plasmodium* species still yield functional phenotypes as demonstrated for *PbΔp52/p36* and *PyΔp52/p36* parasites complemented with PyP52/PbP36 or PbP52/PyP36 [16]. Indeed, the existence of a conserved interaction interface would support the formation of functional chimeric P36-P52 complexes. Interestingly, it appears that P36 and P52 would also possess the ability to form homodimers as supported by wet-lab results (this paper and [37,38]) and AlphaFold structure prediction. However, at this point, it remains unclear to us whether P36-P36 and P52-P52 homodimer formation bears any functional relevance.

Although we could recombinantly obtain a *P. falciparum* P52-P36 fusion protein, its low yields and contamination issues hindered further structural and functional studies. Hence, we describe here a novel “epitope tagging” assay to explore the accessibility of *Plasmodium* proteins to the action of neutralizing antibodies. The assay relies on the genetic modification of the parasite to insert an epitope tag in the protein of interest, and on a functional assay to measure the neutralizing activity of anti-tag antibodies. Using this approach, we demonstrate that the 6-Cys proteins P36 and P52, which are both essential for productive invasion of hepatocytes, are exposed to the inhibitory activity of antibodies that can efficiently block host cell invasion. Interestingly, neutralization of P36 function was more pronounced than with P52, which is consistent with our previous observations that P36 is a key determinant of host receptor usage [16]. Importantly, our data also show that the position of the epitope tag is critical for efficient neutralization, identifying the membrane-distal part of the P36-P52 complex as a vulnerable region, while targeting the membrane-proximal domains has no inhibitory effect. As P36 is possibly physically linked to host receptor usage (CD81 and/or SR-BI) [16], it is thus plausible that the membrane-distal domains of the P36-P52 complex could be exposed for a host cell interaction, which in turn could be blocked by the anti-tag antibody in our experiments. While our antibody-mediated neutralization assay clearly demonstrates that both P36 and P52 are accessible to neutralizing antibodies in culture, we were not able to detect either of the proteins on the SPZ surface (data not shown), even when activated in medium or in the presence of host cells, corroborating previous observations by Arredondo et al. in *P. yoelii* [30]. Both P36 and P52 are localized in the SPZ micronemes, but only P36 has been shown to be secreted in the supernatant of activated parasites [30]. One study reported that P52 can be detected at the surface of gliding *P. berghei* SPZs [31], however this was not confirmed in *P. yoelii* [30]. Along the same line, proteomic studies identified P36 in total but not surface proteomes of SPZs [43,44]. Whether, when and how P36 and P52 translocate to the SPZ surface thus remains unclear. It is possible that, following microneme secretion, P36 and P52 are present only transiently on the parasite surface, and/or in very small amounts, below the detection limit. Alternatively, P36 and/or P52 may exert their function in a soluble form, possibly after cleavage and shedding in the case of P52.

Using a similar C-terminal or N-terminal epitope tagging approach, we failed to block the function of B9, although this protein is also essential for productive invasion. We cannot exclude the possibility that other regions in B9 could be targeted by neutralizing antibodies. In this regard, we attempted to introduce a triple Flag epitope in the middle of B9 (between its propeller and first 6-Cys domains). However, the resulting SPZs showed a complete abrogation of infectivity, reproducing the knockout phenotype and thus precluding functional assays with anti-tag antibodies. Furthermore, we investigated the possible existence of a functional P36-P52-B9 complex required for SPZ infectivity. Such a complex is hypothesized to exist as P36/P52 and B9 mutants display similar phenotypes [16,31–33]. However, the data presented here do not support the existence of a functional tripartite complex. This may also explain why AlphaFold predictions of the *P. falciparum*, *P. berghei*, *P. yoelii* and *P. vivax* P36-P52-B9 complexes display very poor confidence scores.

In conclusion, we set up an assay to measure the accessibility of specific *Plasmodium* proteins to neutralizing antibodies, and identify the SPZ 6-Cys proteins P36 and P52 as vulnerable targets This method could be generalized to other *Plasmodium* antigens, including in other stages of the parasite life cycle, and could be a valuable approach to screen for potential targets of antibody-mediated antimalarial interventions.

## Materials and methods

### Ethics Statement

All animal work was conducted in strict accordance with the Directive 2010/63/EU of the European Parliament and Council on the protection of animals used for scientific purposes. Protocols were approved by the Ethics Committee Charles Darwin N°005 (approval #44204-2023071122181369).

### Experimental animals, parasites and cell lines

Female SWISS mice (6–8 weeks old, from Janvier Labs) were used for all routine parasite infections. Parasite lines were maintained in mice through intraperitoneal injections of frozen parasite stocks and transmitted to *Anopheles stephensi* mosquitoes for experimental purposes. A drop of blood from the tail was collected in 1ml PBS daily and used to monitor the parasitaemia by flow cytometry. *A. stephensi* mosquitoes were reared at 24°C with 80% humidity and permitted to feed on infected mice that were anaesthetized, using standard methods of mosquito infection as previously described [45]. Post-feeding, *P*. *berghei*-infected mosquitoes were kept at 21°C and fed on a 10% sucrose solution. Salivary gland SPZs were collected from infected mosquitoes between 21 and 28 days post-feeding, by hand dissection and homogenisation of isolated salivary glands in complete DMEM (DMEM supplemented with 10% FCS, 1% Penicillin-Streptomycin and 1% L-Glutamine). Mosquitoes infected with GFP-expressing parasites were sorted under a fluorescence microscope prior to dissection. HepG2 cells (ATCC HB-8065) were cultured in DMEM supplemented with 10% FCS, 1% Penicillin-Streptomycin and 1% L-Glutamine, as previously described [46].

### AlphaFold2-based structure prediction

The structural models of SPZ 6Cys protein complexes were predicted using AlphaFold-Multimer [35]. Twenty-five models were predicted per run and the best models underwent a final relaxation step. The models were evaluated based on the following parameters: AlphaFold-Multimer model confidence (a weighted combination of the predicted template modeling (pTM) and interface predicted template modeling (ipTM) scores, 0.8*ipTM + 0.2*pTM) [35], the predicted aligned error (PAE) matrix [36], the local and global predicted local distance difference test (pLDDT) scores [36], the predicted DockQ (pDockQ) values [47] and the normalised discrete optimised protein energy (zDOPE) scores [48]. Protein-protein interactions were analysed using the PISA server [49]. Molecular graphics and analyses were performed with UCSF ChimeraX [50].

### Recombinant production and purification of the PfP52-P36 fusion protein in Sf9 cells

The Bac-to-Bac Expression system (Thermo Fisher Scientific) was used according to the manufacturer’s protocol to produce recombinant baculovirus. In brief, a pFastBac construct containing the PfP52-P36 gene was transformed into DH10Bac competent cells. Blue/white screening was performed to identify colonies containing the recombinant bacmid DNA and the PureLink HiPure Plasmid miniprep kit (Thermo Fischer Scientific) was used to isolate the bacmid DNA. Sf9 cells were transfected with the isolated bacmid DNA using the Expifectamine (Thermo Fisher Scientific) transfection reagent and a P0 recombinant baculovirus stock was produced. This stock underwent two rounds of amplification to obtain a P2 baculovirus stock. Next, P2 was used for the infection of Sf9 cultures. The infected Sf9 cells were cultured in Sf-900 III SFM (Thermo Fisher Scientific) medium in a 27°C shaking incubator. Five days post transfection, the culture medium was harvested by 20 minutes centrifugation at 16000 x g and filtered through a 0,45 μm filter. The fusion protein contains a His-tag and was purified using a two-step purification protocol consisting of IMAC and SEC. First, the sample was loaded onto an equilibrated HisTrap HP 1 mL column (Cytiva) and washed with buffer A (50 mM Tris-HCl, 500 mM NaCl, 20 mM imidazole, pH 8.0). Next, the sample was eluted using a 0 to 100 % gradient of buffer B (50 mM Tris-HCl, 500 mM NaCl, 1 M imidazole, pH 8.0) and peak fractions were collected. Finally, the peak fractions were pooled, concentrated and injected onto the SEC column (Bio-Rad, Enrich 650), 1.5 column volume of SEC buffer (50 mM Tris-HCl, 500 mM NaCl, pH 8.0) was run over the column and peak fractions were collected. Analysis of the fractions was done by SDS-PAGE and Western blot. To check the purity of the sample, total protein was visualized using Coomassie staining. Presence of the protein of interest was detected by western blot using primary mouse anti-His (Bio-Rad) and secondary goat anti-mouse-HRP antibodies (Sigma). Protein concentration was measured using a NanoDrop (Thermo Fisher Scientific) and samples were stored at room temperature.

### Negative stain electron microscopy

Samples for negative stain electron microscopy (EM) were prepared by incubating 3 μL of P52-P36 fusion protein (0.519 mg/mL undiluted, 1/10 and 1/50 diluted in buffer 50 mM Tris, 150 mM NaCl pH 8) on glow-discharged copper grids (carbon coated, 300 mesh, EM Resolutions, Staffordshire, United Kingdom) for 1 min at room temperature. Grids were washed twice with 20 μL of buffer 25 mM HEPES-NaOH, 200 mM NaCl, 0.5 mM TCEP pH 7.4 and twice with 20 μL of deionized water. The sample was stained with 5 μL of 2 % uranyl acetate solution for 30 s. Excess staining solution was removed with filter paper and the grids were air dried. Negative stain images were acquired at 120 kV using a Tecnai™ G2 Spirit Bio TWIN microscope (Thermo Fisher Scientific, Eindhoven, The Netherlands).

### Small-angle X-ray scattering

Small-angle X-ray scattering (SAXS) was conducted at the BioSAXS beamline SWING (SOLEIL, Gif-sur-Yvette, France) [51]. SEC-SAXS data were acquired on a Shodex KW404-4F column, pre-equilibrated in 20 mM HEPES, 200 mM NaCl, 3% glycerol, pH 8. A sample of 55 µL (4.2 mg/mL) was injected and subsequently eluted at a flow rate of 0.3 mL/min. Scattering data were collected with an exposure time of 990 msec and a dead time of 10 msec, calibrated to absolute units using the scattering of pure water [52]. Data processing and analysis was performed using the ATSAS package and BioXTAS RAW [53–55]. As the SEC-SAXS profile contained convoluted peaks of distinct scattering particles, the data were processed using Evolving Factor Analysis (EFA) to extract the scattering profile of the individual components [56]. The information on data collection and derived structural parameters is summarized in **Supplementary Table 1**. Molecular models of the *P. falciparum* P52-P36 fusion protein were generated using AlphaFold [36,57] Theoretical scattering curves of the AlphaFold models and their respective fits to the experimental data were calculated using FoXS [58]. SAXS-based ensemble modelling was carried out using BILBOMD [59,60]. BilboMD Classic was used to generate conformational ensembles by sampling six Rg bins, with minimal and maximal Rg values set at 7% and 35% of the experimentally determined Rg, respectively. For each Rg bin, 800 conformations were generated, yielding 4800 conformers in the run from which a minimal ensemble was selected. The overall goodness-of-fit between the final models and the experimental data are reported through the calculation of a χ2 value, with Nk being the number of points, σ(qj) the standard deviations, and c a scaling factor.

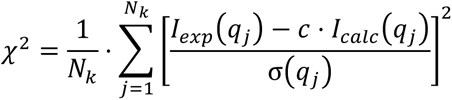

Furthermore, comparisons of theoretical and experimental scattering curves include the presentation of a residuals plot (Δ/σ vs. q, where Δ indicates the difference between experimental and calculated intensities), which enables a local inspection of the model fit to the data.

### Generation of P36-FlagC, P52-V5C/P36-FlagC, FlagN-B9 and B9-FlagInt parasites

#### Plasmid constructs

Genetic complementation of Δ*p36* [16], Δ*p52p36* [16] and Δ*b9* [32] parasites was achieved by double crossover homologous recombination using a vector containing a hDHFR cassette and a 3’ homology arm corresponding to the 5’ sequence of the HSP70 promoter of the GFP cassette present in the parental lines. To generate the construct for C-terminal tagging of P36, we first inserted a fragment consisting of the *p36* promoter, an *Xho*I site, a 3xFlag epitope and the *p36* 3’ UTR, between *Kpn*I and *Eco*RI sites of the vector. This insert was provided as a synthetic gene (Eurofins Genomics). In a second step, the coding sequence of P36 was cloned into the *Xho*I site, resulting in the final construct used to transfect Δ*p36* parasites to generate the P36-FlagC line. This plasmid was further modified by sequential insertion the p52 promoter and the intergenic sequence containing *p52* 3’ UTR and *p36* promoter, and the p52 coding sequence containing a 3xFlag epitope inserted towards the C-terminus upstream of the predicted omega site. This construct was used to transfect the Δ*p52p36* parasites to generate the P52-V5C/P36-FlagC parasite line. For tagging of B9, the ΔpropΔ6cys1 plasmid [32] was modified by inserting two fragments corresponding to the N-terminal portion of B9 containing a 3xFlag epitope (amplified by PCR from a synthetic gene) and the second exon of b9 (amplified by PCR from parasite genomic DNA), resulting in the construct used for N-terminal tagging of B9 in the FlagN-B9 line. To generate the B9-FlagInt parasites, two fragments corresponding to the propeller domain and the 6-cys domains were cloned into the ΔpropΔ6cys1 plasmid, with introduction of a 3xFlag between the two fragments. All cloning steps were performed using the CloneAmp HiFi PCR premix and the In-Fusion HD Cloning Kit (Takara). The plasmids were verified by Sanger DNA sequencing (Eurofins Genomics) and linearized with *Kpn*I and *Nhe*I before transfection. All the primers used for plasmid assembly are listed in **Supplementary Table S2**.

#### Transfection and selection

For parasite transfection, schizonts were purified from overnight cultures of Δ*p36,* Δ*p52p36* or Δ*b9 P. berghei* parasite lines [16,32], transfected with 10 μg of linearized constructs by electroporation using the AMAXA Nucleofector device (program U033), as previously described [61], and then immediately injected intravenously into the tail vein of SWISS mice. To permit the selection of resistant transgenic parasites, pyrimethamine (35 mg/L) was added to the drinking water and administered to mice, starting one day after transfection. The mice were monitored daily by flow cytometry to detect the reappearance of parasites. When parasitemia reached at least 1%, mouse blood was collected for preparation of frozen stocks and isolation of parasites for genomic DNA extraction and genotyping.

### Generation of P52-FlagC, FlagN-P52 and FlagN-P36 parasites using CRISPR-Cas9

#### sgRNA guide plasmids

Using the Chop-Chop (https://chopchop.cbu.uib.no) and Benchling (https://www.benchling.com) program, two 20 bp guide RNA sequences were selected upstream of PAM motifs in the 5’ and 3’ ends of *p36* and *p52* coding sequence. Their complementary oligonucleotides were subsequently designed and optimized in the Takara Primer design tool (https://www.takarabio.com/learning-centers/cloning/primer-design-and-other-tools). A guanosine nucleotide was added at the 5’ end of the forward oligonucleotide for enhancing transcriptional initiation [39]. Paired oligonucleotides were annealed and cloned into *Bsm*BI and *Bsa*I sites of the psgRNA_Pf-Pb U6_2targets plasmid using the In-Fusion HD Cloning Kit (Takara), resulting in the insertion of the guide RNA immediately downstream of PfU6 or PbU6 promoter, respectively [39]. This plasmid contains a *hDHFR*-*yfcu* cassette, for positive selection by pyrimethamine and negative selection by 5-fluorocytosine (5-FC) [62,63]. The resulting sgRNA guide plasmids were checked by Sanger sequencing prior to transfection.

#### Donor DNA templates for DNA repair by double homologous recombination

The donor templates to modify *p36* or *p52* locus were provided as synthetic genes, containing a 5’ upstream fragment serving for 5’ homologous recombination, a coding sequence containing a 3xFlag, and a 3’ downstream fragment serving for 3’ homologous recombination. Synthetic genes were flanked by two *Xho*I sites, allowing linearization prior to transfection. The oligonucleotide and synthetic gene sequences are listed in **Supplementary Table S2**.

#### Parasite transfection and selection

For parasite transfection, schizonts were purified from an overnight culture of the parent parasite line PbCasDiCre-GFP [40] and transfected with a mix of 10 μg of sgRNA plasmid and 10 μg of linearized donor template by electroporation using the AMAXA Nucleofector device (program U033), as previously described [61], and immediately injected intravenously into the tail vein of SWISS mice. To permit the selection of resistant transgenic parasites, pyrimethamine (35 mg/L) was added to the drinking water and administered to mice, starting one day after transfection and for a total of 4-5 days. Following withdrawal of pyrimethamine, the mice were monitored daily by flow cytometry to detect the reappearance of parasites. When parasitemia reached at least 1%, mouse blood was collected for preparation of frozen stocks and isolation of parasites for genomic DNA extraction and genotyping.

### Generation of triple transgenic PbPfP52P36B9

PbPfP52P36B9 parasites were generated using CRISPR-Cas9 in three successive transfection steps. For the three genes, we only modified the 6-Cys and/or propeller domains, which are presumably essential for complex formation and/or function, while preserving the *P. berghei* N-terminal signal peptide and C-terminal GPI anchor sequences, to ensure proper expression and trafficking of the modified proteins. Two 20 bp guide RNA sequences were selected upstream of PAM motifs in the 5’ and 3’ ends of *p36*, *p52* or *b9* coding sequence, and paired oligonucleotides were cloned into the psgRNA_Pf-Pb U6_2targets plasmid, as described above. The donor templates to modify *p36, p52* or *b9* locus were provided as synthetic genes, containing a 5’ upstream fragment serving for 5’ homologous recombination, a sequence encoding PfP52 (residues 44-340), PfP36 (residues 80-379) or PfB9 (residues 29-833), and a 3’ downstream fragment serving for 3’ homologous recombination. Parasite transfection and selection was performed as described above for N-terminal tagging of P36 and 52. The resulting PbPfP52P36B9 triple transgenic parasites were genotyped by PCR. The oligonucleotide and synthetic gene sequences are listed in **Supplementary Table S2**.

### Genotyping PCR

Blood collected from infected mice was passed through a CF11 column (Whatman) to deplete leucocytes. The RBCs collected were then centrifuged and lysed with 0.2% saponin (Sigma) to recover parasite material for genomic DNA isolation using a kit (Qiagen DNA Easy Blood and Tissue Kit), according to the manufacturer’s instructions. Genomic DNA served as template for PCR, using specific primer combinations designed to detect the wild-type or recombined loci. All PCR reactions were carried out using Recombinant Taq DNA Polymerase (5U/μl from Thermo Scientific) and standard PCR cycling conditions. All the primer sequences are listed in **Supplementary Table S2**.

### *In vitro* infection assays

HepG2 and HepG2/CD81 cells [64] were seeded in collagen-coated culture plates, at a density of 30,000 cells/well in a 96-well plate for flow cytometry analysis and immunofluorescence assays, 24 hours prior to infection with SPZs. On the day of infection, the culture medium in the wells was refreshed with complete DMEM, followed by the addition of 10,000 SPZs/well for flow cytometry analysis or 1,000 SPZs/well for immunofluorescence assays. For SPZ invasion inhibition assays, anti-Flag (Low Endotoxin, Azide-Free purified rat IgG2a, clone L5, Biolegend, #637328), anti-V5 (Azide-free rabbit IgG, EPR12989, Abcam, #ab250573) or an irrelevant isotype control antibody (Low Endotoxin, Azide-Free purified rat IgG2a isotype control, #400565) were added at various concentrations in the culture medium together with the SPZs. Infected cultures were incubated for 3 hours at 37°C. The wells were then washed twice with complete DMEM and then incubated for another 24–48 hours at 37°C before analysis by flow cytometry. For quantification of infected cells by flow cytometry, the cultures were trypsinized after two washes with PBS, followed by addition of complete DMEM and one round of centrifugation. After discarding the supernatant, the cells were directly resuspended in FACS buffer (PBS + 1% FCS) and analyzed on a Guava EasyCyte 6/2L bench cytometer equipped with 488 nm and 532 nm lasers (Millipore).

### Immunofluorescence assays

For immunofluorescence assays on HepG2 and HepG2/CD81 infected cultures, the cells were washed twice with PBS, then fixed with 4% PFA for 10 minutes followed by two washes with PBS, quenching with 0.1 M glycine for 5 minutes, permeabilization with 1% Triton X-100 for 5 minutes before washes with PBS and blocking in PBS with 3% bovine serum albumin (BSA). Cells were then incubated for 1h with goat anti-UIS4 primary antibody (1:500, Sicgen), followed by Alexa Fluor 594- or Alexa Fluor 488-conjugated donkey anti-goat secondary antibodies (1:1000, Life Technologies). Nuclei were stained with Hoechst 33342 (Life Technologies). Samples were then imaged on a Zeiss Axio Observer.Z1 fluorescence microscope equipped with a LD Plan-Neofluar 40x/0.6 Corr Ph2 M27 objective. The same exposure conditions were maintained for all the conditions in order to allow comparisons.

Images were processed with ImageJ for adjustment of contrast. For immunofluorescence analysis of Flag-tagged parasites, SPZs collected from infected mosquito salivary glands were deposited on poly-L-lysine coated coverslips, fixed with 4% PFA and permeabilized with 1% Triton X-100. Parasites were labelled with anti-Flag mouse antibodies (M2 clone, Sigma) and anti-TRAP rabbit antibodies [65], AlexaFluor 594-conjugated secondary antibodies (Life Technologies). Nuclei were stained with Hoechst 33342. Coverslips were mounted on glass slides with ProLong™ Diamond Antifade Mountant (Life Technologies), and imaged on a Zeiss Axio Observer.Z1 fluorescence microscope equipped with a LD Plan-Neofluar 40x/0.6 Corr Ph2 M27 objective. Images were processed with ImageJ for adjustment of contrast.

### Ultrastructure expansion microscopy

SPZs were collected from infected mosquito salivary glands, centrifuged at 3800 g during 4 min at 4 °C and resuspended in 1X PBS. In some experiments, microneme secretion was stimulated by incubation for 15 min at 37°C in a buffer containing 1% BSA and 1% ethanol, as described [66,67]. Parasites were sedimented on poly-D-lysine coverslips (100 μL/coverslip) during 30 min at room temperature (RT). Samples were then prepared for U-ExM as previously described [66]. Briefly, coverslips were incubated overnight in a 2% Acrylamide/1.4% Formaldehyde solution at 37 °C. Gelation was then performed in 10% ammonium persulfate (APS)/10% Temed in monomer solution (19% Sodium Acrylate; 10% Acrylamide; 0.1% BIS-Acrylamide in PBS) during 1 h at 37 °C. Following gelation, denaturation was performed in 200mM SDS, 200mM NaCl and 50mM Tris pH 9.0 during 90 min at 95 °C. A first round of expansion was performed by incubating the gels thrice in ultrapure water for 30 min at RT. Gels were then washed in PBS twice for 15 min to remove excess water and blocked with 2% BSA in PBS for 30 min at RT. Staining was then performed by incubation with primary antibodies diluted at 1/250 in PBS containing 2% BSA at RT overnight with 120–160 rpm shaking. We used antibodies against Flag (clone L5, Biolegend), V5 (SV5-Pk1, ThermoFisher Scientific) or TRAP [65]. The next day, gels were washed 3 times for 10 min in PBS-Tween 0.1%. Incubation with the secondary antibodies was performed for 3 h at RT with 120–160 rpm shaking, followed by 3 washes of 10 minutes in PBS-Tween 0.1%. Directly after washing, gels were expanded for a second round in ultrapure water for 30 min, thrice. For imaging, 5 mm x 5 mm gel pieces were cut from the expanded gels and mounted between glass slides and Poly-D-Lysine coated coverslips. Acquisitions were made on a Zeiss LSM700 confocal microscope, using the ZEN 2012 SP5 FP3 (black) version 14.0.0.0 (Zeiss). Images were processed with ImageJ for adjustment of contrast.

### Statistical analysis

Statistical significance in the *in vitro* infection assays was assessed by two-way ratio paired t tests. All statistical tests were computed with GraphPad Prism 10 (GraphPad Software). *In vitro* experiments were performed with a minimum of three technical replicates per experiment. Quantitative source data are provided in **Supplementary Table S3**.

## Acknowledgements

We thank Thierry Houpert and Louise Deltour-Foglio for rearing of mosquitoes, Freddy Frischknecht and Jessica Kehrer for the kind gift of anti-TRAP antibodies, and Wai-Hong Tham and Melanie Dietrich for sharing their Pf52 construct with us. This work was funded by grants from the Laboratoire d’Excellence ParaFrap (ANR-11-LABX-0024), the Agence Nationale de la Recherche (ANR-20-CE18-0013 and ANR-22-CE18-0007) and the Fondation pour la Recherche Médicale (EQU201903007823). ML was supported by a ‘DIM 1Health’ doctoral fellowship awarded by the Conseil Régional d’Ile-de- France. LDV is a doctoral fellow supported by the FWO-Vlaanderen (11P4B24N). LB was supported by a Christian Boulin fellowship to learn how to work with the Sf21, Sf9 and Hi5 insect cell lines for recombinant protein production in Heidelberg at the EMBL Facility (we explicitly wish to acknowledge Kim Remans, Karine Lapouge, Yexin Xie and Jacob Scheurich). The authors wish to thank the staff or the SWING beam line at the SOLEIL synchrotron (Aurélien THUREAU) for outstanding beam line support. The authors acknowledge use of the CalcUA and VSC supercomputing facilities and wish to thank the staff for outstanding support.

## Supplementary information

**Supplementary Figure S1.**
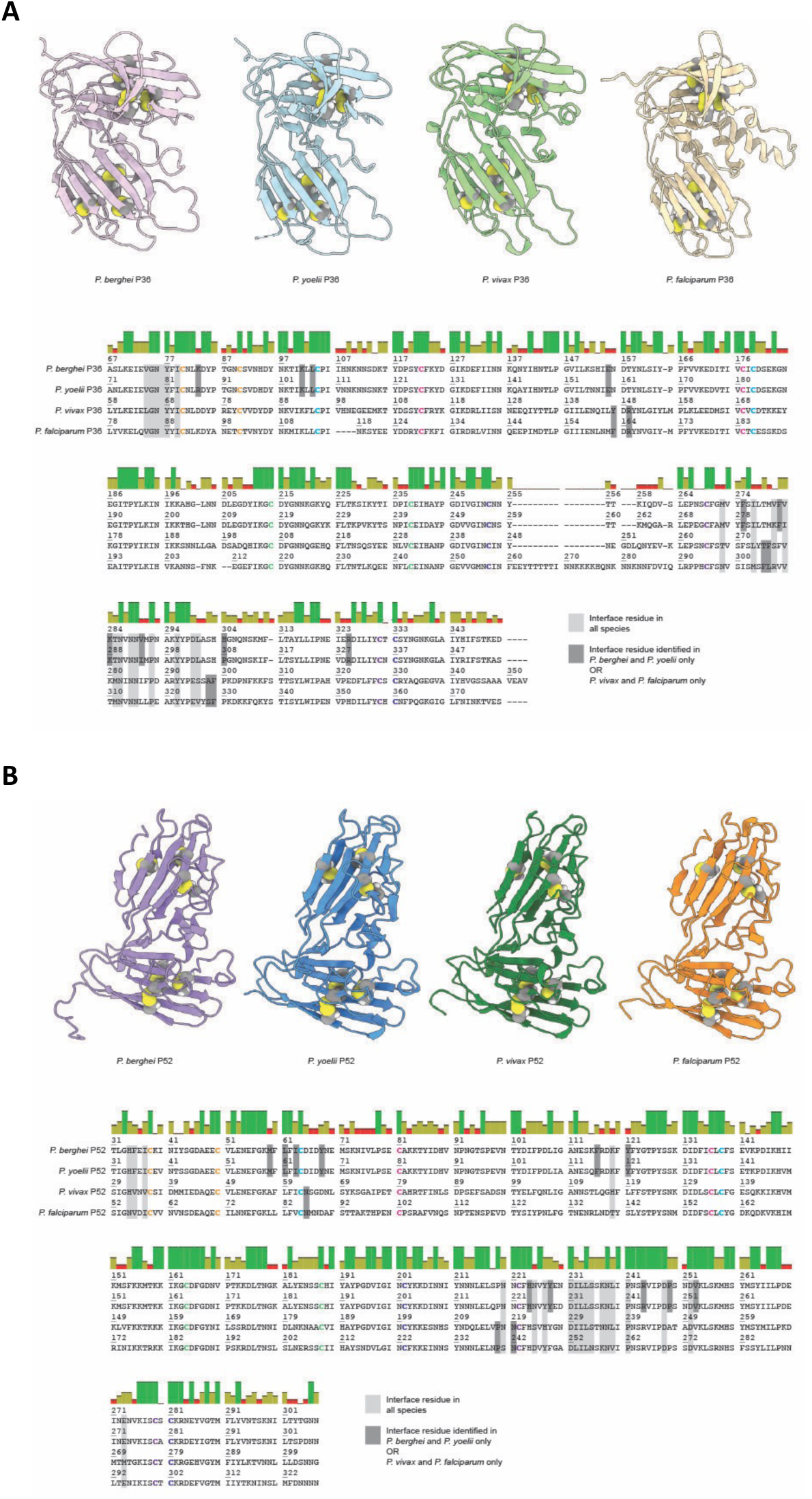
Conservation of P36 and P52 structures, sequences, and interface residues. **A**. The top section shows cartoon representations of P36 AlphaFold models color-coded as in Figure 2. Disulfides are shown as spheres and colored by heteroatom. The bottom section displays a multiple sequence alignment (MSA) of P36 sequences from *P. berghei*, *P. yoelii*, *P. vivax*, and *P. falciparum*. Cysteines involved in the formation of an intramolecular disulfide are indicated in bold. Interface residues in all species are indicated by the light grey areas, whereas those only found in i) both *P. berghei* and *P. yoelii* or ii) both *P. vivax* and *P. falciparum* are highlighted by dark grey areas. The colored bars above the MSA represent the percentage of sequence identity: green (100%), green-brown (between 30% and 100%), and red (below 30%). **B**. Same analysis for P52.

**Supplementary Figure S2.**
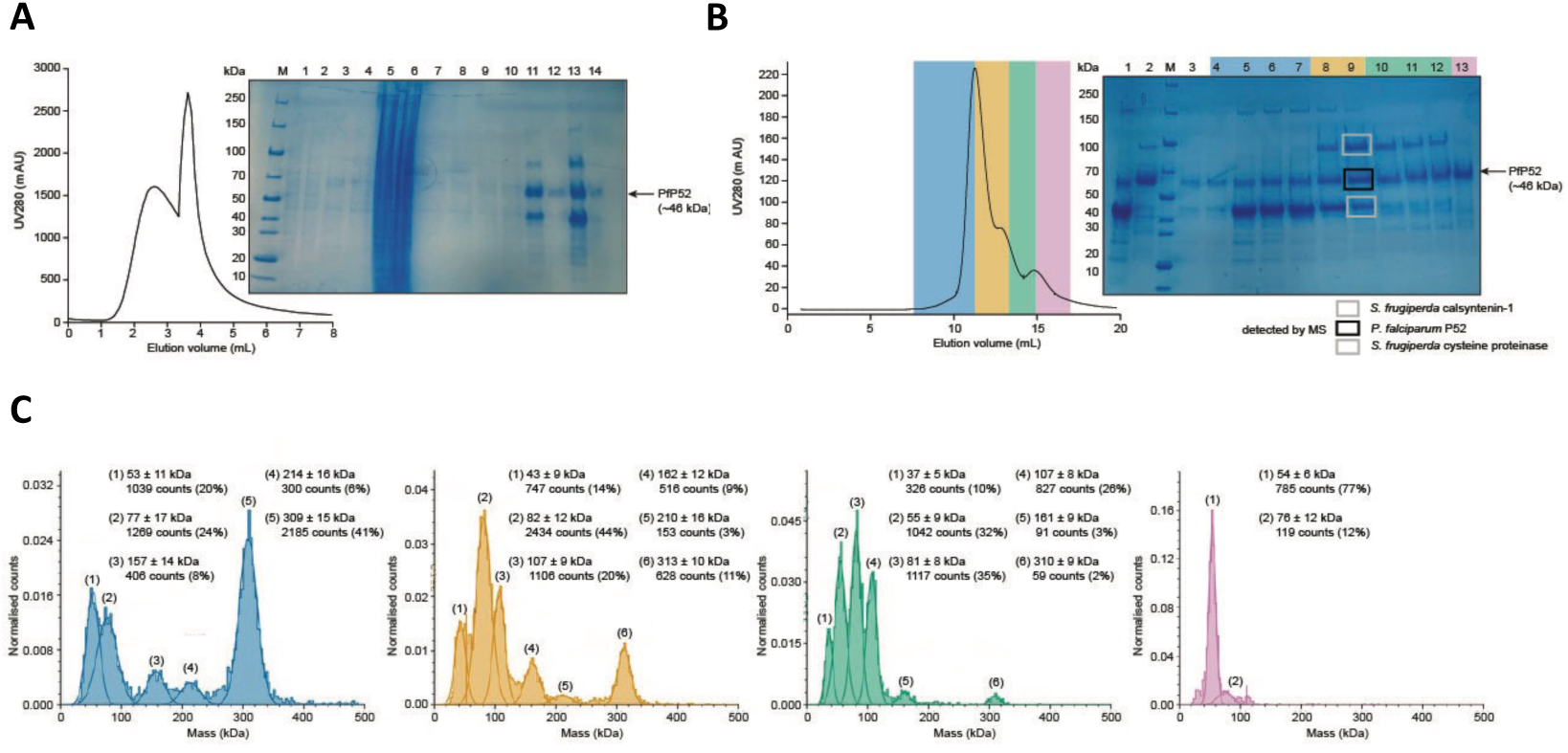
Recombinant production and purification trials for *P. falciparum* P52. **A**. IMAC elution profile after having loaded the insect cell culture supernatant onto the column (Complete His-tag 1 mL, Roche). The elution was performed in a single step with 100% elution buffer and was paused after ∼3 mL for ∼5 minutes to incubate the column with the elution buffer to accelerate the process, as evidenced by the sudden increase in UV280 absorbance when elution is resumed. The inset shows an SDS-PAGE analysis of various analyzed fractions. Two IMACs were performed with samples from two cell cultures, grown and transfected in parallel. *Lane 1, whole cell culture 1 before harvesting. Lane 2, whole cell culture 2 before harvesting. Lane 3, supernatant of centrifuged cell culture 1. Lane 4, supernatant of centrifuged cell culture 2. Lane 5, pellet of cell culture 1. Lane 6, pellet of cell culture 2. Lane 7, filtered supernatant of cell culture 1 loaded onto the column. Lane 8, filtered supernatant of cell culture 2 loaded onto the column. Lane 9, flow through of culture 1 after loading sample. Lane 10, flow through of culture 2 after loading sample. Lane 11, pooled elution fractions of IMAC 1. Lane 12, pre-elution fraction of IMAC 1. Lane 13, pooled elution fractions of IMAC 2. Lane 14, pre-elution fraction of IMAC 2. Lane M, PageRuler Unstained Broad Range Protein Ladder.* The black arrow indicates the expected molecular mass for recombinant *P. falciparum* P52 (PfP52, ∼46 kDa). **B**. SEC elution profile on pooled fractions after IMAC (fractions 2A and 2B). The volume of the pooled fractions was ∼12 mL, two consecutive SEC runs were performed (Superdex 200 10/30 increase, Cytiva). The inset shows an SDS-PAGE analysis of various analyzed fractions from one of the two identical runs*. Lanes 1 and 2, fractions from the other SEC run. Lane 3, fraction 1. Lanes 4 to 7, fractions 2 until 5, which fall under the area highlighted in blue. Lanes 8 and 9, fractions 6 and 7, which fall under the area highlighted in orange. Lanes 10 to 12, fractions 8 until 10, which fall under the area highlighted in green. Lane 13, fraction 11, highlighted in purple. Lane M, PageRuler Unstained Broad Range Protein Ladder.* SEC fractions were pooled for subsequent mass photometry analysis. These pools are color-coded as follows: pool 1 (fractions 4 to 7, blue), pool 2 (fractions 8 to 9, yellow), pool 3 (fractions 10 to 12, green), and pool 4 (fraction 13, purple). The boxes indicated excised gel bands that were analyzed by mass spectrometry (MS) to confirm the identity of *P. falciparum* P52 (black box) and insect cell contaminants grey boxes). **C**. Mass photometry data collected for the various SEC pools, which shows the heterogeneity in P52 monomer vs. oligomer species.

**Supplementary Figure S3.**
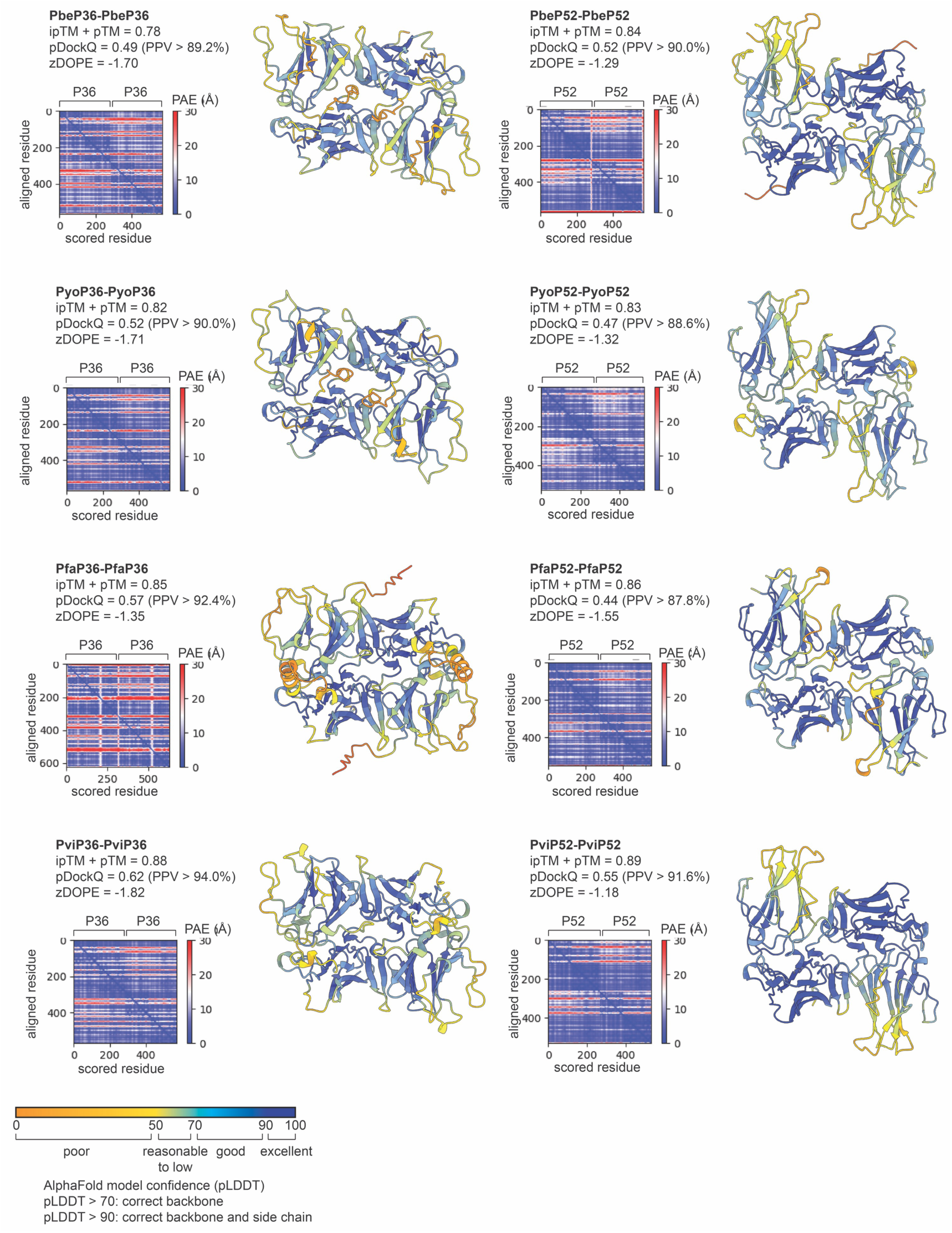
Overview of the AlphaFold2 prediction models for the P36-P36 and P52-P52 homodimers and associated quality metrics. The models are displayed as cartoon representations and are colored according to the predicted local distance difference test (pLDDT) score, which reflects (local) model quality as indicated by the legend at the bottom. For all structures, the predicted aligned error (PAE), the normalized discrete optimized protein energy (zDOPE), the pDockQ and AlphaFold-Multimer model confidence (0.8*ipTM + 0.2*pTM) are also shown.

**Supplementary Figure S4.**
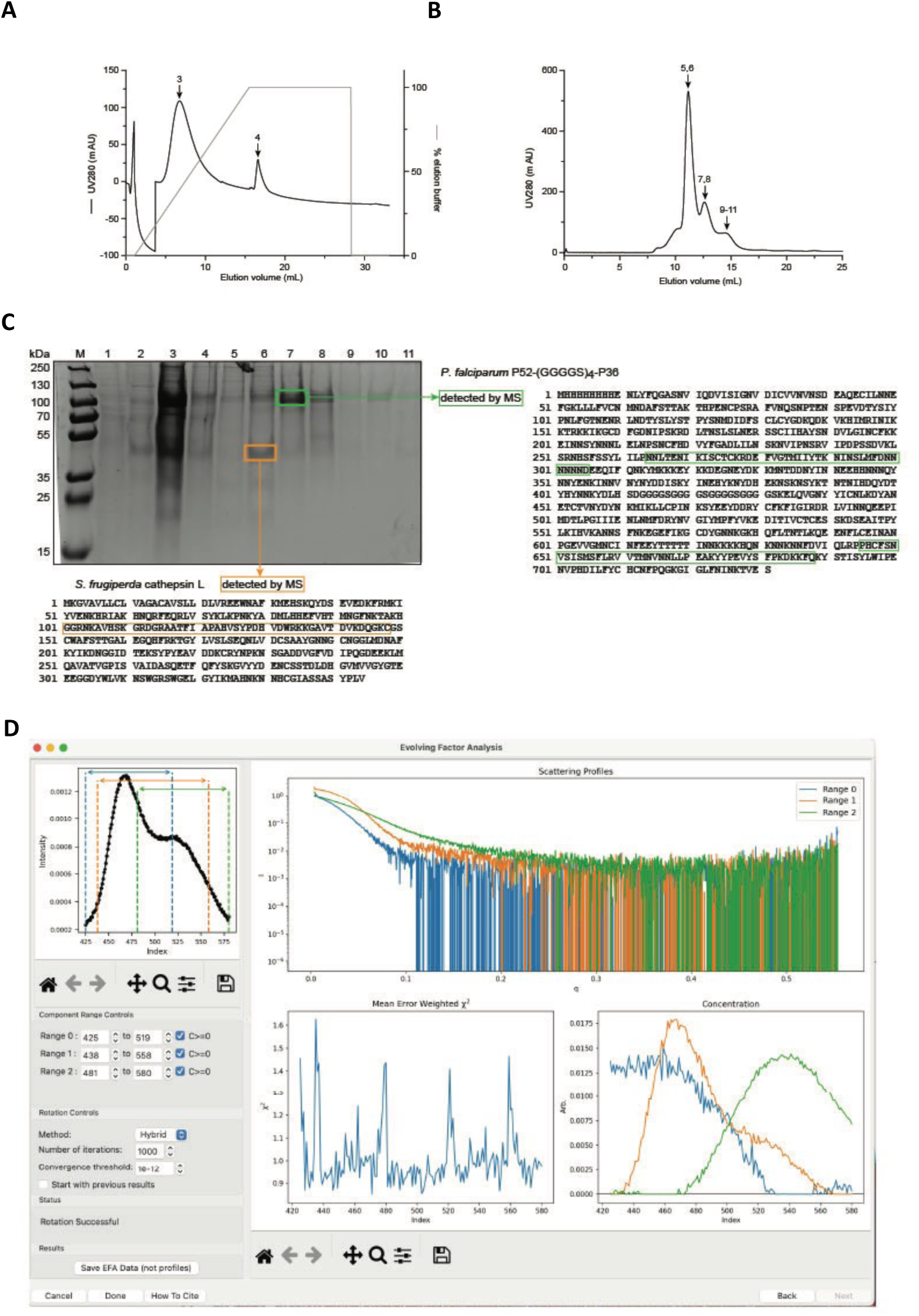
Recombinant production and purification for the *P. falciparum* P52-P36 fusion protein. **A**. IMAC elution profile after having loaded the insect cell culture supernatant onto the column (HisTrap HP 1 mL, Cytiva). The elution was performed through an elution gradient. **B**. SEC elution profile on pooled fractions after IMAC (fractions 2 to 6). **C**. SDS-PAGE analysis of the fractions collected during SEC and IMAC. *Lane 1, filtered supernatant of cell culture 1 loaded onto the column. Lane 2, wash fraction prior to elution. Lane 3, IMAC elution peak. Lane 4, IMAC elution peak. Lanes 5 to 11, SEC elution fractions. Lane M, PageRuler Plus PreStained Protein Ladder.* The colored boxes indicate the excised gel bands that were analyzed via MS to confirm the identity of the *P. falciparum* P52-P36 fusion construct (∼83 kDa) and the insect cell contaminant (∼38 kDa). **D**. Screenshot of the SAXS data processing employing evolving factor analysis to deconvolute the elution peaks obtained during SEC-SAXS to obtain the scattering curve for the *P. falciparum* P52-P36 fusion construct as presented in Figure 3. The evolving factor analysis identifies three scattering components, of which the third component (colored green) corresponds to the *P. falciparum* P52-P36 fusion construct.

**Supplementary figure S5.**
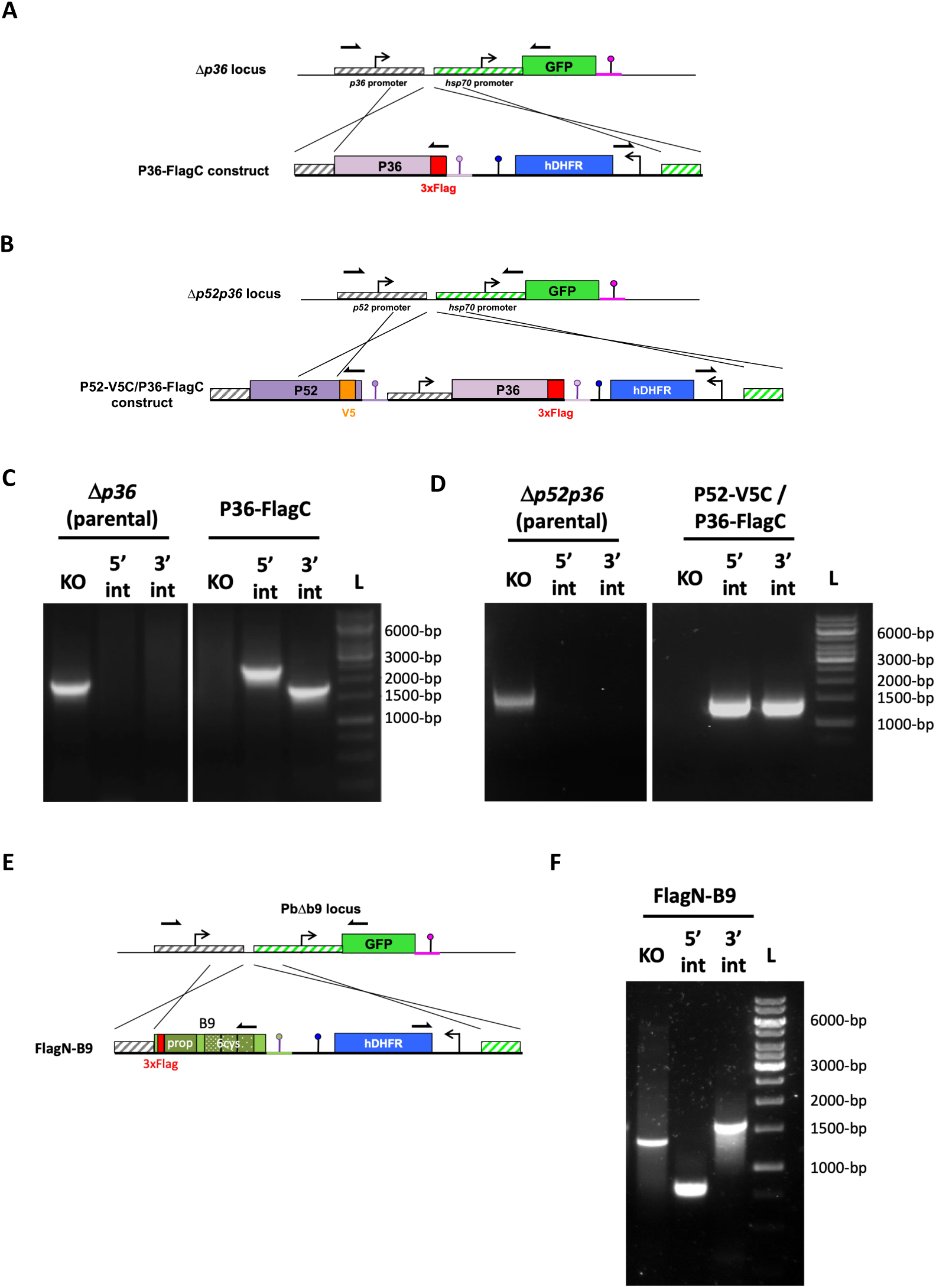
Genetic tagging of 6-Cys proteins in *P. berghei*. **A**. Genetic strategy to generate the P36-FlagC parasites, based on genetic complementation of Δ*p36* with a construct harboring a P36 coding sequence with a C-terminal 3xFlag in addition to a hDHFR pyrimethamine resistance cassette. **B**. Genetic strategy to generate the P52-V5C/P36-FlagC parasites, based on genetic complementation of Δ*p52p36* with a construct harboring a P52 coding sequence with a C-terminal V5 epitope, a P36 coding sequence with a C-terminal 3xFlag, and the hDHFR pyrimethamine resistance cassette. **C-D**. Genotyping of P36-FlagC (C) and P52-V5C/P36-FlagC (D) parasites by PCR using primers combinations specific for the parental genome (WT) or for the 5’ and 3’ recombination events. **E**. Genetic strategy to generate the FlagN-B9 parasites, based on genetic complementation of GFP-expressing Δ*b9* with a construct harboring a B9 coding sequence with a N-terminal 3xFlag epitope in addition to a hDHFR pyrimethamine resistance cassette. **F**. Genotyping of FlagN-B9 parasites by PCR using primers combinations specific for the parental genome (WT) or for the 5’ and 3’ recombination events.

**Supplementary Figure S6.**
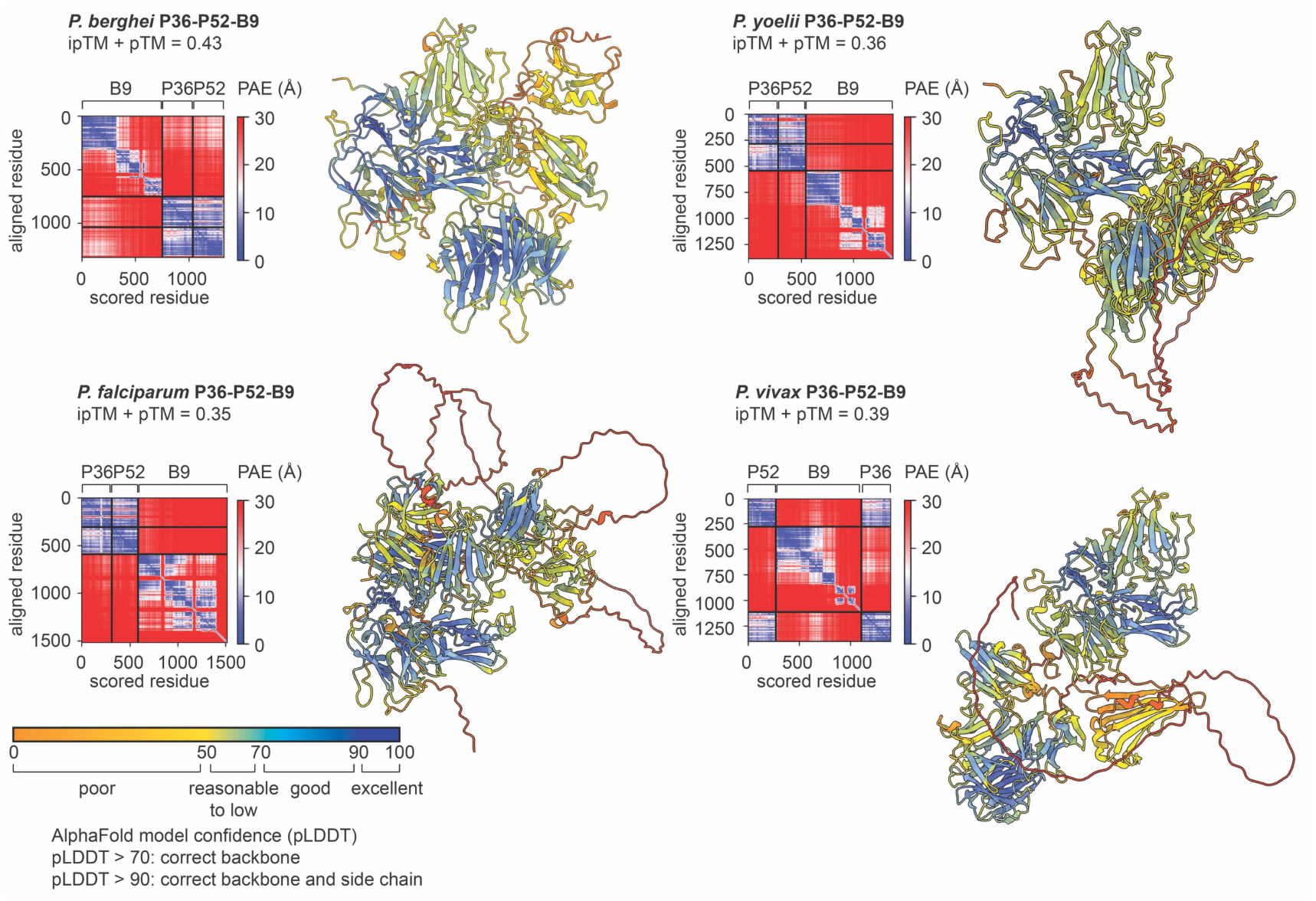
AlphaFold prediction models for the P36-P52-B9 complex. The heterotrimer models of P36-P52-B9 in *P. berghei*, *P. yoelii*, *P. vivax*, and *P. falciparum* are displayed as cartoon representations and are colored according to the predicted local distance difference test (pLDDT) score, which reflects (local) model quality as indicated by the legend at the bottom. For all structures, the predicted aligned error (PAE) and AlphaFold-Multimer model confidence (0.8*ipTM + 0.2*pTM) are also shown.

**Supplementary Figure S7.**
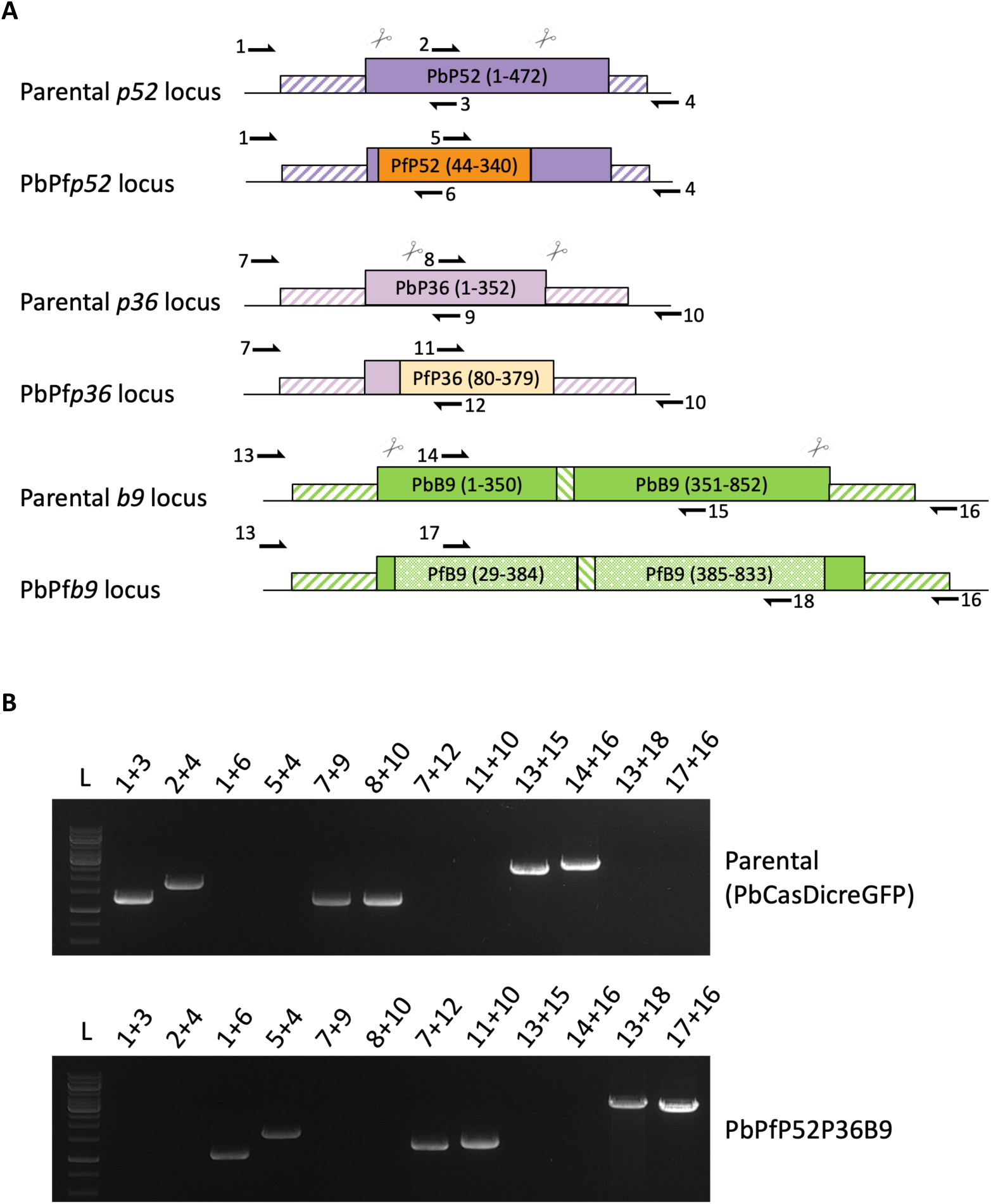
Generation of transgenic PbPfP36P52B9 parasites. **A**. Strategy to replace PbP52, PbP36 and PbB9 in a PbCasDiCreGFP parasite line using CRISPR to successively generate falciparumized PbPfP52, PbPfP36P52 and PbPfP36P52B9 lines. The PbCasDiCreGFP parasites were co-transfected with a linearized DNA repair construct synthetically designed with a plasmid encoding gene specific sgRNA guides and a pyrimethamine-resistance cassette (hDHFR). **B**. PCR analysis of the genomic DNA obtained from the parental PbCasDiCreGFP line and the recombinant line PbPfP52P36B9. Confirmation of the expected recombination events was achieved with primer combinations specific for *P. berghei* or *P. falciparum* gene sequences.

## References

1. Datoo MS, Natama MH, Somé A, Traoré O, Rouamba T, Bellamy D, et al. Efficacy of a low-dose candidate malaria vaccine, R21 in adjuvant Matrix-M, with seasonal administration to children in Burkina Faso: a randomised controlled trial. The Lancet. 2021;397. doi:10.1016/S0140-6736(21)00943-0

2. Datoo MS, Natama HM, Somé A, Bellamy D, Traoré O, Rouamba T, et al. Efficacy and immunogenicity of R21/Matrix-M vaccine against clinical malaria after 2 years’ follow-up in children in Burkina Faso: a phase 1/2b randomised controlled trial. Lancet Infect Dis. 2022;22. doi:10.1016/S1473-3099(22)00442-X

3. Olotu A, Fegan G, Wambua J, Nyangweso G, Leach A, Lievens M, et al. Seven-Year Efficacy of RTS,S/AS01 Malaria Vaccine among Young African Children. New England Journal of Medicine. 2016;374: 2519–2529. doi:10.1056/NEJMoa1515257

4. 4. RTSS Clinical Trials Partnership. Efficacy and safety of RTS,S/AS01 malaria vaccine with or without a booster dose in infants and children in Africa: final results of a phase 3, individually randomised, controlled trial. The Lancet. 2015;386: 31–45. doi:10.1016/S0140-6736(15)60721-8

5. White MT, Verity R, Griffin JT, Asante KP, Owusu-Agyei S, Greenwood B, et al. Immunogenicity of the RTS,S/AS01 malaria vaccine and implications for duration of vaccine efficacy: secondary analysis of data from a phase 3 randomised controlled trial. Lancet Infect Dis. 2015;15: 1450–8. doi:10.1016/S1473-3099(15)00239-X

6. Miura K, Flores-Garcia Y, Long CA, Zavala F. Vaccines and monoclonal antibodies: new tools for malaria control. Clin Microbiol Rev. 2024;37. doi:10.1128/cmr.00071-23

7. Kayentao K, Ongoiba A, Preston AC, Healy SA, Doumbo S, Doumtabe D, et al. Safety and Efficacy of a Monoclonal Antibody against Malaria in Mali. New England Journal of Medicine. 2022;387. doi:10.1056/nejmoa2206966

8. Gaudinski MR, Berkowitz NM, Idris AH, Coates EE, Holman LA, Mendoza F, et al. A Monoclonal Antibody for Malaria Prevention. New England Journal of Medicine. 2021;385: 803–814. doi:10.1056/NEJMoa2034031

9. Wu RL, Idris AH, Berkowitz NM, Happe M, Gaudinski MR, Buettner C, et al. Low-Dose Subcutaneous or Intravenous Monoclonal Antibody to Prevent Malaria. New England Journal of Medicine. 2022;387: 397–407. doi:10.1056/NEJMoa2203067

10. Lyke KE, Berry AA, Mason K, Idris AH, O’Callahan M, Happe M, et al. Low-dose intravenous and subcutaneous CIS43LS monoclonal antibody for protection against malaria (VRC 612 Part C): a phase 1, adaptive trial. Lancet Infect Dis. 2023;23: 578–588. doi:10.1016/S1473-3099(22)00793-9

11. Aliprandini E, Tavares J, Panatieri RH, Thiberge S, Yamamoto MM, Silvie O, et al. Cytotoxic anti-circumsporozoite antibodies target malaria sporozoites in the host skin. Nature Microbiology. 2018. pp. 1224–1233. doi:10.1038/s41564-018-0254-z

12. Stewart MJ, Nawrot RJ, Schulman S, Vanderberg JP. Plasmodium berghei sporozoite invasion is blocked in vitro by sporozoite-immobilizing antibodies. Infect Immun. 1986;51: 859–864. doi:10.1128/iai.51.3.859-864.1986

13. Aguirre-Botero MC, Wang LT, Formaglio P, Aliprandini E, Thiberge JM, Schön A, et al. Cytotoxicity of human antibodies targeting the circumsporozoite protein is amplified by 3D substrate and correlates with protection. Cell Rep. 2023;42. doi:10.1016/j.celrep.2023.112681

14. Aguirre-Botero MC, Pacios O, Celli S, Aliprandini E, Gladston A, Thiberge J-M, et al. Late killing of Plasmodium berghei sporozoites in the liver by an anti-circumsporozoite protein antibody. Elife. 2025;14. doi:10.7554/eLife.105291

15. Silvie O, Rubinstein E, Franetich JF, Prenant M, Belnoue E, Rénia L, et al. Hepatocyte CD81 is required for Plasmodium falciparum and Plasmodium yoelii sporozoite infectivity. Nat Med. 2003;9: 93–96. doi:10.1038/nm808

16. Manzoni G, Marinach C, Topçu S, Briquet S, Grand M, Tolle M, et al. Plasmodium P36 determines host cell receptor usage during sporozoite invasion. Elife. 2017;6: e25903. doi:10.7554/eLife.25903

17. Lyons FMT, Gabriela M, Tham WH, Dietrich MH. Plasmodium 6-Cysteine Proteins: Functional Diversity, Transmission-Blocking Antibodies and Structural Scaffolds. Frontiers in Cellular and Infection Microbiology. 2022. doi:10.3389/fcimb.2022.945924

18. Gerloff DL, Creasey A, Maslau S, Carter R. Structural models for the protein family characterized by gamete surface protein Pfs230 of Plasmodium falciparum. Proc Natl Acad Sci U S A. 2005;102. doi:10.1073/pnas.0502378102

19. Tonkin ML, Arredondo SA, Loveless BC, Serpa JJ, Makepeace KAT, Sundar N, et al. Structural and biochemical characterization of Plasmodium falciparum 12 (Pf12) reveals a unique interdomain organization and the potential for an antiparallel arrangement with Pf41. Journal of Biological Chemistry. 2013;288. doi:10.1074/jbc.M113.455667

20. Higgins MK, Carrington M. Sequence variation and structural conservation allows development of novel function and immune evasion in parasite surface protein families. Protein Science. 2014. doi:10.1002/pro.2428

21. Templeton TJ, Iyer LM, Anantharaman V, Enomoto S, Abrahante JE, Subramanian GM, et al. Comparative analysis of apicomplexa and genomic diversity in eukaryotes. Genome Res. 2004;14. doi:10.1101/gr.2615304

22. Aravind L, Iyer LM, Wellems TE, Miller LH. Plasmodium Biology: Genomic Gleanings. Cell. 2003. doi:10.1016/S0092-8674(03)01023-7

23. Arredondo SA, Kappe SHI. The s48/45 six-cysteine proteins: mediators of interaction throughout the Plasmodium life cycle. International Journal for Parasitology. 2017. pp. 409–423. doi:10.1016/j.ijpara.2016.10.002

24. Arredondo SA, Schepis A, Reynolds L, Kappe SHI. Secretory Organelle Function in the Plasmodium Sporozoite. Trends in Parasitology. 2021. doi:10.1016/j.pt.2021.01.008

25. Loubens M, Vincensini L, Fernandes P, Briquet S, Marinach C, Silvie O. Plasmodium sporozoites on the move: Switching from cell traversal to productive invasion of hepatocytes. Mol Microbiol. 2021;115: 870–881. doi:10.1111/mmi.14645

26. Kumar N, Wizel B. Further characterization of interactions between gamete surface antigens of Plasmodium falciparum. Mol Biochem Parasitol. 1992;53. doi:10.1016/0166-6851(92)90013-A

27. Dietrich MH, Chmielewski J, Chan L-J, Tan LL, Adair A, Lyons FMT, et al. Cryo-EM structure of endogenous *Plasmodium falciparum* Pfs230 and Pfs48/45 fertilization complex. Science (1979). 2025;389. doi:10.1126/science.ady0241

28. Taechalertpaisarn T, Crosnier C, Bartholdson SJ, Hodder AN, Thompson J, Bustamante LY, et al. Biochemical and functional analysis of two plasmodium falciparum blood-stage 6-Cys proteins: P12 and P41. PLoS One. 2012;7: e41937. doi:10.1371/journal.pone.0041937

29. Dietrich MH, Chan LJ, Adair A, Boulet C, O’Neill MT, Tan LL, et al. Structure of the Pf12 and Pf41 heterodimeric complex of Plasmodium falciparum 6-cysteine proteins. FEMS Microbes. 2022;3. doi:10.1093/femsmc/xtac005

30. Arredondo SA, Swearingen KE, Martinson T, Steel R, Dankwa DA, Harupa A, et al. The Micronemal Plasmodium Proteins P36 and P52 Act in Concert to Establish the Replication-Permissive Compartment Within Infected Hepatocytes. 2018;8: 413. doi:10.3389/fcimb.2018.00413

31. Ishino T, Chinzei Y, Yuda M. Two proteins with 6-cys motifs are required for malarial parasites to commit to infection of the hepatocyte. Mol Microbiol. 2005;58: 1264–1275. doi:10.1111/j.1365-2958.2005.04801.x

32. Fernandes P, Loubens M, Marinach C, Coppée R, Baron L, Grand M, et al. Plasmodium sporozoites require the protein B9 to invade hepatocytes. iScience. 2023. doi:10.1016/j.isci.2023.106056

33. Annoura T, Van Schaijk BCL, Ploemen IHJ, Sajid M, Lin JW, Vos MW, et al. Two Plasmodium 6-Cys family-related proteins have distinct and critical roles in liver-stage development. FASEB Journal. 2014;28: 2158–2170. doi:10.1096/fj.13-241570

34. van Schaijk BCL, Janse CJ, van Gemert GJ, van Dijk MR, Gego A, Franetich JF, et al. Gene discruption of Plasmodium falcifarum p52 results in attenuation of malaria liver stage development in cultured primary human hepatocytes. PLoS One. 2008;3: e3549. doi:10.1371/journal.pone.0003549

35. 35. Evans R, O’Neill M, Pritzel A, Antropova N, Senior A, Green T, et al. Protein complex prediction with AlphaFold-Multimer. bioRxiv. 2022.

36. Jumper J, Evans R, Pritzel A, Green T, Figurnov M, Ronneberger O, et al. Highly accurate protein structure prediction with AlphaFold. Nature. 2021;596: 583–589. doi:10.1038/s41586-021-03819-2

37. Kaushansky A, Douglass AN, Arang N, Vigdorovich V, Dambrauskas N, Kain HS, et al. Malaria parasites target the hepatocyte receptor EphA2 for successful host infection. Science (1979). 2015;350: 1089–1092. doi:10.1126/science.aad3318

38. Vigdorovich V, Patel H, Watson A, Raappana A, Reynolds L, Selman W, et al. Coimmunization with Preerythrocytic Antigens alongside Circumsporozoite Protein Can Enhance Sterile Protection against *Plasmodium* Sporozoite Infection. Microbiol Spectr. 2023;11. doi:10.1128/spectrum.03791-22

39. Shinzawa N, Nishi T, Hiyoshi F, Motooka D, Yuda M, Iwanaga S. Improvement of CRISPR/Cas9 system by transfecting Cas9-expressing Plasmodium berghei with linear donor template. Commun Biol. 2020;3. doi:10.1038/s42003-020-01138-2

40. Das S, Unhale T, Marinach C, Valeriano Alegria B del C, Roux C, Madry H, et al. Constitutive expression of Cas9 and rapamycin-inducible Cre recombinase facilitates conditional genome editing in Plasmodium berghei. Sci Rep. 2025;15: 2949. doi:10.1038/s41598-025-87114-4

41. Mueller AK, Camargo N, Kaiser K, Andorfer C, Frevert U, Matuschewski K, et al. Plasmodium liver stage developmental arrest by depletion of a protein at the parasite-host interface. Proc Natl Acad Sci U S A. 2005;102: 3022–3027. doi:10.1073/pnas.0408442102

42. Vanderberg JP, Frevert U. Intravital microscopy demonstrating antibody-mediated immobilisation of Plasmodium berghei sporozoites injected into skin by mosquitoes. Int J Parasitol. 2004;34: 991–996.

43. Lindner SE, Swearingen KE, Harupa A, Vaughan AM, Sinnis P, Moritz RL, et al. Total and putative surface proteomics of malaria parasite salivary gland sporozoites. Mol Cell Proteomics. 2013;12: 1127–1143. doi:10.1074/mcp.M112.024505

44. Swearingen KE, Lindner SE, Shi L, Shears MJ, Harupa A, Hopp CS, et al. Interrogating the Plasmodium Sporozoite Surface: Identification of Surface-Exposed Proteins and Demonstration of Glycosylation on CSP and TRAP by Mass Spectrometry-Based Proteomics. Yates J, editor. PLoS Pathog. 2016;12: e1005606. doi:10.1371/journal.ppat.1005606

45. Ramakrishnan C, Delves MJ, Lal K, Blagborough AM, Butcher G, Baker KW, et al. Laboratory maintenance of rodent malaria parasites. Methods Mol Biol. 2013;923: 51–72. doi:10.1007/978-1-62703-026-7_5

46. Silvie O, Franetich JF, Boucheix C, Rubinstein E, Mazier D. Alternative invasion pathways for plasmodium berghei sporozoites. Int J Parasitol. 2007;37: 173–182. doi:10.1016/j.ijpara.2006.10.005

47. Bryant P, Pozzati G, Elofsson A. Improved prediction of protein-protein interactions using AlphaFold2. Nat Commun. 2022;13. doi:10.1038/s41467-022-28865-w

48. Shen M, Sali A. Statistical potential for assessment and prediction of protein structures. Protein Science. 2006;15. doi:10.1110/ps.062416606

49. Krissinel E, Henrick K. Inference of Macromolecular Assemblies from Crystalline State. J Mol Biol. 2007;372. doi:10.1016/j.jmb.2007.05.022

50. Meng EC, Goddard TD, Pettersen EF, Couch GS, Pearson ZJ, Morris JH, et al. UCSF ChimeraX: Tools for structure building and analysis. Protein Science. 2023;32. doi:10.1002/pro.4792

51. Thureau A, Roblin P, Pérez J. BioSAXS on the SWING beamline at Synchrotron SOLEIL. J Appl Crystallogr. 2021;54. doi:10.1107/S1600576721008736

52. Orthaber D, Bergmann A, Glatter O. SAXS experiments on absolute scale with Kratky systems using water as a secondary standard. J Appl Crystallogr. 2000;33. doi:10.1107/S0021889899015216

53. Panjkovich A, Svergun DI. CHROMIXS: Automatic and interactive analysis of chromatography-coupled small-angle X-ray scattering data. Bioinformatics. 2018;34. doi:10.1093/bioinformatics/btx846

54. Manalastas-Cantos K, Konarev P V., Hajizadeh NR, Kikhney AG, Petoukhov M V., Molodenskiy DS, et al. ATSAS 3.0: Expanded functionality and new tools for small-angle scattering data analysis. J Appl Crystallogr. 2021;54. doi:10.1107/S1600576720013412

55. Hopkins JB. BioXTAS RAW 2: new developments for a free open-source program for small-angle scattering data reduction and analysis. J Appl Crystallogr. 2024;57. doi:10.1107/S1600576723011019

56. Meisburger SP, Taylor AB, Khan CA, Zhang S, Fitzpatrick PF, Ando N. Domain Movements upon Activation of Phenylalanine Hydroxylase Characterized by Crystallography and Chromatography-Coupled Small-Angle X-ray Scattering. J Am Chem Soc. 2016;138: 6506–6516. doi:10.1021/jacs.6b01563

57. Tunyasuvunakool K, Adler J, Wu Z, Green T, Zielinski M, Žídek A, et al. Highly accurate protein structure prediction for the human proteome. Nature. 2021;596. doi:10.1038/s41586-021-03828-1

58. Schneidman-Duhovny D, Hammel M, Sali A. FoXS: a web server for rapid computation and fitting of SAXS profiles. Nucleic Acids Res. 2010;38. doi:10.1093/nar/gkq461

59. Schneidman-Duhovny D, Hammel M, Tainer JA, Sali A. FoXS, FoXSDock and MultiFoXS: Single-state and multi-state structural modeling of proteins and their complexes based on SAXS profiles. Nucleic Acids Res. 2016;44. doi:10.1093/nar/gkw389

60. Pelikan M, Hura GL, Hammel M. Structure and flexibility within proteins as identified through small angle X-ray scattering. Gen Physiol Biophys. 2009;28. doi:10.4149/gpb_2009_02_174

61. Janse CJ, Ramesar J, Waters AP. High-efficiency transfection and drug selection of genetically transformed blood stages of the rodent malaria parasite Plasmodium berghei. Nat Protoc. 2006;1: 346–356.

62. Braks JAM, Franke-Fayard B, Kroeze H, Janse CJ, Waters AP. Development and application of a positive-negative selectable marker system for use in reverse genetics in Plasmodium. Nucleic Acids Res. 2006;34. doi:10.1093/NAR/GNJ033

63. Manzoni G, Briquet S, Risco-Castillo V. A rapid and robust selection procedure for generating drug-selectable marker-free recombinant malaria parasites. Sci Rep. 2014;99210: 1–10. doi:10.1038/srep04760

64. Silvie O, Greco C, Franetich JF, Dubart-Kupperschmitt A, Hannoun L, van Gemert GJ, et al. Expression of human CD81 differently affects host cell susceptibility to malaria sporozoites depending on the Plasmodium species. Cell Microbiol. 2006;8: 1134–1146. doi:10.1111/j.1462-5822.2006.00697.x

65. Klug D, Goellner S, Kehrer J, Sattler J, Strauss L, Singer M, et al. Evolutionarily distant I domains can functionally replace the essential ligandbinding domain of plasmodium trap. Elife. 2020;9: e57572. doi:10.7554/eLife.57572

66. Loubens M, Marinach C, Paquereau C-E, Hamada S, Hoareau-Coudert B, Akbar D, et al. The claudin-like apicomplexan microneme protein is required for gliding motility and infectivity of Plasmodium sporozoites. PLoS Pathog. 2023;19: e1011261. doi:10.1371/journal.ppat.1011261

67. Brown KM, Sibley LD, Lourido S. High-Throughput Measurement of Microneme Secretion in Toxoplasma gondii. Methods in Molecular Biology. 2020. pp. 157–169. doi:10.1007/978-1-4939-9857-9_9

