## Supplemental Table S1 for "A structure-based epitope tagging approach identifies vulnerable sites on the malarial P36-P52 protein complex for antibody-mediated neutralization of *Plasmodium* sporozoites"

**Table S1.** SAXS data collection and scattering-derived parameters for *P. falciparum* P52-P36.

| **Data collection parameters** | |
|---|---|
| Beam line | SWING (PROXIMA) |
| Wavelength (Å) | 0.99 |
| $q$ range ($Å^{-1}$)* | 0.0005 - 0.5543 |
| Concentration (mg $ml^{-1}$) (mode) | 4.2 (SEC-SAXS) |
| Buffer conditions | 20 mM HEPES, 200 mM NaCl, 3% glycerol, pH 8.0 |
| Temperature (°C) | 20 |
| **Structural parameters**§ | |
| *I(0)* ($cm^{-1}$) [from Guinier] | 0.85 |
| $R_g$ (Å) [from Guinier] | 41.18 |
| *I(0)* ($cm^{-1}$) [from *p(r)*] | 0.87 |
| $R_g$ (Å) [from *p(r)*] | 45.90 |
| E.R. | 2.06 |
| $D_{max}$ (Å) | ~210 |
| Porod volume estimate, $V_p$ ($Å^3$) | 179,887 |
| Porod exponent | 3.0 |
| **Molecular mass determination** | |
| MM (kDa) [from *SAXSMoW* on final merged curve] | 83.1 ($q$ = 0.20 $Å^{-1}$) |
| MM (kDa) [from $Q_R$ on final merged curve] | 82.9 ($q$ = 0.15 $Å^{-1}$) |
| MM (kDa) [from $V_p$/1.7] | 105.8 |
| Calculated MM from sequence (kDa) | 84.4 |
| **Ensemble modeling** | |
| Conformer generation and selection | BILBOMD |
| Protein regions selected to be rigid/fixed | 19-289, 432-730 |
| **Software employed** | |
| Data processing and analysis | ATSAS, BioXTAS RAW |
| **SASBDB** | |
| **Entry** | SASDY36 |

Abbreviations: *I(0)*, extrapolated scattering intensity at zero angle; $R_g$, radius of gyration calculated using either Guinier approximation (from Guinier) or the indirect Fourier transform package GNOM [from *p(r)*]; *MM*, molecular mass; $D_{max}$, maximal particle dimension; $V_p$, Porod volume; E.R., elongation ratio

*Momentum transfer $|q| = 4\pi\sin(\theta)/\lambda$
